# Perceptual learning of random acoustic patterns: Impact of temporal regularity and attention

**DOI:** 10.1101/2023.03.13.532336

**Authors:** Hanna Ringer, Erich Schröger, Sabine Grimm

**Affiliations:** International Max Planck Research School on Neuroscience of Communication (IMPRS NeuroCom), Max Planck Institute for Human Cognitive and Brain Sciences, Leipzig, Germany; Cognitive and Biological Psychology, Wilhelm Wundt Institute for Psychology, Leipzig University, Leipzig, Germany; Research Group Neurocognition of Music and Language, Max Planck Institute for Empirical Aesthetics, Frankfurt am Main, Germany; Physics of Cognition Lab, Institute of Physics, Chemnitz University of Technology, Chemnitz, Germany

**Keywords:** perceptual learning, auditory memory, acoustic patterns, temporal regularity, attention, EEG

## Abstract

Perceptual learning is a powerful mechanism to enhance perceptual abilities and to form robust memory representations of previously unfamiliar sounds. Memory formation through repeated exposure takes place even for random and complex acoustic patterns devoid of semantic content. The current study sought to scrutinise how perceptual learning of random acoustic patterns is shaped by two potential modulators: temporal regularity of pattern repetition and listeners’ attention. To this end, we adapted an established implicit learning paradigm and presented short acoustic sequences that could contain embedded repetitions of a certain sound segment (i.e., pattern) or not. During each experimental block, one repeating pattern recurred across multiple trials, while the other patterns were presented in only one trial. During the presentation of sound sequences that contained either temporally regular or jittered within-trial pattern repetitions, participants’ attention was directed either towards or away from the auditory stimulation. Overall, we found a memory-related modulation of the event-related potential (ERP) and an increase in inter-trial phase coherence for patterns that recurred across multiple trials (compared to non-recurring patterns), accompanied by a performance increase in a (within-trial) repetition detection task when listeners attended the sounds. Remarkably, we show a memory-related ERP effect even for the first pattern occurrence per sequence when participants attended the sounds, but not when they were engaged in a visual distractor task. These findings suggest that learning of unfamiliar sound patterns is robust against temporal irregularity and inattention, but attention facilitates access to established memory representations upon first occurrence within a sequence.

## Introduction

Perceptual learning is a powerful mechanism to enhance perception and acquire novel memory representations throughout one’s lifetime. It is defined as the experience-dependent gain in perceptual capacity through growing experience with a certain, usually previously unfamiliar, type of stimulus material (Gibson, 1969; Gilbert et al., 2001; Irvine et al., 2000). In the auditory modality, repeated exposure to a novel class of sounds rapidly improves listeners’ abilities to perceptually parse them and, for instance, efficiently discriminate different exemplars (Wright & Zhang, 2009).

This plays a particularly crucial role in challenging listening situation in which different stimulus exemplars are hard to distinguish because they are highly similar or the signal quality is suboptimal (Banai & Lavie, 2020; Irvine, 2018; Samuel & Kraljic, 2009). In turn, sharpened perception of relevant sensory input through experience and efficient recognition of already known information facilitates successful interaction and adaptive behaviour in challenging auditory environments (Bregman, 1990; Winkler et al., 2009). For instance, repeated exposure not only allows listeners to become more sensitive to stimulus details that they could not retain upon first hearing (McDermott et al., 2013) and segregate different sound sources more readily (Woods & McDermott, 2018), but also supports higher-level cognitive functions including speech and music perception (Irvine et al., 2000; Kraus & Banai, 2007; Norris et al., 2003).

The current study focuses on a specific aspect of perceptual learning, that is the formation of item-specific memories for particular (random) stimulus exemplars (out of a pool of perceptually highly similar stimuli) through repeated exposure. Other than an overall, item-unspecific increase in perceptual capacity as a result of growing experience with a certain type of stimulus material, this should be reflected in a selective enhancement of perceptual performance for stimulus exemplars that recur across trials compared to others that do not recur over trials. While there is compelling evidence that the human auditory system has a remarkable capacity to form robust memories even of random and meaningless acoustic patterns (as will be reviewed below), research on how implicit perceptual learning of such patterns is modulated by different aspects of the learning context remains rather scarce. The present study investigates the influence of two potential modulators that are widely discussed to enhance perceptual and cognitive processing across a variety of different contexts: temporal regularity of the sensory input and attention to the stimulation (and, in particular, relevant features therein).

### Behavioural and electrophysiological markers of acoustic pattern learning

Most studies investigating memory formation for random acoustic patterns used the so-called implicit noise-learning paradigm (Agus et al., 2010), which was initially established with white noise as stimulus material, but later applied to different types of random acoustic patterns (e.g., Agus & Pressnitzer, 2021; Bianco et al., 2020; Kang et al., 2017; Ringer et al., 2022a). In their original experiment, Agus and colleagues (2010) presented listeners with white noise stimuli that either consisted of two seamless presentations of the same segment or of random noise and asked them to decide in every trial whether or not the sound contained a repetition. Unbeknownst to the participants, one specific segment recurred across multiple trials within an experimental block while all other patterns occurred in only one trial. The authors argued that the gradual increase of hit rate in the repetition detection task for recurring “reference” noise patterns (relative to other non-recurring patterns) indicates successful memory formation for reference patterns, which in turn improved repetition detection (Agus et al., 2010). Learning was characterised as implicit, since the repetitions of the reference pattern across trials happened unbeknownst to the listeners, and it occurred fast after just a few presentations of the reference pattern and despite interference of irrelevant noises in between. Acquired memory representations proved to be long-lasting and robust against temporal and spectral transformation, i.e., the performance benefit persisted over two weeks and even for time-compressed or -reversed versions of the reference patten (Agus et al., 2010).

This finding was replicated and extended by multiple subsequent studies. Robust memory effects through repeated exposure to a reference pattern were found not only for white noise as stimulus material (Agus et al., 2010; Agus & Pressnitzer, 2013; Andrillon et al., 2015, 2017; Dauer et al., 2022; Song & Luo, 2017; Viswanathan et al., 2016), but also for “correlated” noise (Ringer et al., 2022a), “tone clouds” of varying complexity (Agus & Pressnitzer, 2021; Kumar et al., 2014), sequences of short tones (Bianco et al., 2020; Herrmann et al., 2021), and temporal patterns of clicks (Kang et al., 2017, 2018, 2021). Successful memory formation was not dependent on immediate within-sound repetition (Agus & Pressnitzer, 2013; Ringer et al., 2022a) and occurred even with much less frequent repetitions of the reference pattern (i.e., only every three minutes on average; Bianco et al., 2020). A recent study showed that, beyond the sensitivity benefit in a perceptual task, repeated exposure enabled above-chance behavioural recognition of learnt reference patterns in a subsequent surprise two-alternative forced-choice memory test, suggesting at least some degree of active accessibility of the memory representations (Ringer et al., 2022a).

The memory-related behavioural changes were accompanied by characteristic neural signatures, as measured using several different neural markers and neuroimaging methods. A magnetoencephalography (MEG) study reported stronger inter-trial phase coherence (ITPC) of low-frequency oscillations (3-8 Hz) for recurring reference (compared to other) white noise patterns that gradually increased over the course of a block along with behavioural sensitivity in the repetition detection task (Luo et al., 2013). Remarkably, different reference patterns elicited distinguishable neural phase patterns (Luo et al., 2013) and distinguishable spatial activation patterns in planum temporale and hippocampus in a functional magnetic resonance imaging (fMRI) study (Kumar et al., 2014). It was argued that repeated exposure to the reference pattern increased perceptual sensitivity to subtle acoustic features via resetting the phase of ongoing low-frequency neural oscillations to these features (Luo et al., 2013). A similar increase in ITPC of low-frequency oscillations for reference patterns was also found using electroencephalography (EEG; Andrillon et al., 2015, 2017). A study using sequences of short tones as stimuli reported an earlier emergence, followed by a reduction in magnitude, of sustained neural activity for sequences that contained repetitions of recurring reference compared to novel patterns (Herrmann et al., 2021). Another line of research investigated event-related potentials (ERPs) relative to the repeating onsets of single pattern repetitions within a continuous auditory sequence. These studies showed a prominent fronto-central negativity that peaked around 200 to 300 ms after pattern onset, sometimes preceded by an earlier frontal positivity (Andrillon et al., 2015, 2017; Hodapp & Grimm, 2021). The overall shape of the waveform was reminiscent of the “noise-locked” negativity reported in response to the onset of the repeating sound segment in periodic white noise sequences (Berti et al., 2000; Kaernbach et al., 1998; Ringer et al., 2022b), and, critically, its amplitude was modulated by the recurrence of a certain reference pattern across multiple trials (Andrillon et al., 2015, 2017; Hodapp & Grimm, 2021). Notably, a difference between learnt reference patterns and other patterns emerged already after the first pattern presentation per trial, i.e., before any within-trial repetition, likely reflecting longer-term memory formation across trials (Andrillon et al., 2015).

### Temporal regularity and attention as potential modulators of perceptual learning

While there is compelling evidence for perceptual learning of different types of random acoustic patterns through repeated exposure, less is known about how different aspects of the learning context shape memory formation. Temporal regularity of the stimulation and participants’ attentional focus are commonly known to facilitate perception (for reviews, see, e.g., Chun & Turk-Browne, 2007; Henry & Herrmann, 2014). Thus, it is plausible to assume that these two factors also influence implicit perceptual learning of random acoustic patterns, which recent research began to investigate.

Temporal regularity of auditory stimuli, inherently linked to enhanced predictability, was demonstrated to reliably benefit perceptual performance across a variety of tasks, including discrimination tasks related to different stimulus features (Barnes & Jones, 2000; Geiser et al., 2012; Jones et al., 2002, 2006), concurrent sound stream segregation (Andreou et al., 2011; Sohoglu & Chait, 2016), detection of near-threshold stimuli (Lawrance et al., 2014) and stimulus changes (Chang et al., 2019; Henry & Obleser, 2012), and repetition detection within continuous sounds (Rajendran et al., 2016). Dynamic Attending Theory postulates rhythmic fluctuations of attention and perceptual sensitivity that align with temporally regular stimulus sequences (Henry & Herrmann, 2014; Jones, 1976, 2019; Large & Jones, 1999) via entrainment of low-frequency neural oscillations to the external stimulation (Calderone et al., 2014; Lakatos et al., 2008, 2019), such that peaks in neural excitability coincide with critical time windows (Lakatos et al., 2005, 2009; Schroeder & Lakatos, 2009). A conducive effect of temporal regularity in the stimulation was also found for memory formation (Hanslmayr et al., 2019; Hickey & Race, 2021). Two recent studies that used the implicit learning paradigm to investigate perceptual learning of acoustic patterns suggested that temporal regularity of pattern repetitions is not a prerequisite for successful memory formation, but might have facilitative impact (Dauer et al., 2022; Hodapp & Grimm, 2021). In the EEG, a modulation of the pattern-related negativity was observed for reference tone patterns, equally in sequences with isochronous and temporally jittered pattern repetitions (Hodapp & Grimm, 2021). Conversely, Dauer and colleagues (2022) showed that behavioural learning was reduced by onset uncertainty and by temporal irregularity of white noise pattern repetitions. Interestingly, this disadvantage was ameliorated as soon as some predictable and isochronous sequences were included in the experimental block (Dauer et al., 2022).

Likewise, attention and memory formation are closely interdependent. Attention acts as a selection mechanism to allocate limited processing resources to particularly relevant sensory input and to enhance its perceptual representation, which in turn paves the way for the formation of robust longer-term memories (Chun & Turk-Browne, 2007). In fact, there is some evidence that attention is a necessary requirement for perceptual learning: For instance, perception of noise-vocoded speech only improved when participants directed their attention to the speech stimuli, but not when they focussed on another auditory or visual task during a training phase (Huyck & Johnsrude, 2012). Similarly, understanding and subsequent recognition of degraded speech required attention, while it was unimpaired by a shift of attention away from the speech stimuli for clear speech (Wild et al., 2012). However, there is also a large body of research that demonstrated successful memory formation for novel sounds in the absence of attention, suggesting that the influence of attention during encoding of the to-be-learnt stimuli depends on perceptual difficulty and is stronger for explicit than for implicit learning (Chun & Turk-Browne, 2007). Even in the absence of attention to the auditory stimulation, the emergence of temporal regularities in continuous acoustic sequences can be detected, as reflected in an amplitude increase of the sustained response (Barascud et al., 2016; Southwell et al., 2017; Southwell & Chait, 2018), and exploited to segregate overlapping sound streams (Masutomi et al., 2016). Also, Berti and colleagues (2000) showed comparable ERP waveforms relative to the onset of the repeating segment within periodic white noise sequences both while participants performed an auditory task or read a book, i.e., with and without attention to the stimulation (Berti et al., 2000). While repetition detection as such does not require memories of the repeating pattern across multiple trials, it does rely on a successful sensory memory comparison within the trial and can be considered at least a preliminary stage of perceptual learning on a shorter time scale. Using the implicit learning paradigm, two studies probed memory formation across trials for specific white noise patterns during inattention (Andrillon et al., 2015, 2017). They found evidence for successful perceptual learning of reference patterns, indexed by a significant amplitude increase of pattern-related ERPs relative to other patterns, while listeners performed a distractor task (Andrillon et al., 2015) and even during rapid eye movement (REM) sleep (Andrillon et al., 2017), suggesting an automatic, pre-attentive nature of the memory formation process. However, to date no study has directly compared the memory effect between different levels of attention towards the auditory stimulation and, in particular, the pattern repetitions therein. Thus, although substantial perceptual learning for random acoustic patterns occurred in the absence of attention to the auditory stimulation, it remains unclear whether attention improves implicit memory formation.

### The present study

The goal of the present EEG study is to scrutinise the effect of two potential modulators on perceptual learning of random acoustic patterns: temporal regularity of pattern repetitions within auditory sequences and listeners’ attention. Previous studies investigated only either of these two factors in isolation, but not their interaction. Also, they mostly only probed the memory effect at separate levels of the factor (e.g., with an auditory task that focussed listeners’ attention on the pattern repetitions *or* with a visual distractor task that directed their attention away from the auditory stimulation). To test whether temporal regularity and attention actually *modulate* the memory effect, i.e., whether the memory effect differs significantly in magnitude between factor levels, in the present study we manipulated both factors in a two-by-two within-subject design. We used an adapted version of the established implicit learning paradigm (Agus et al., 2010) and presented to-be-learnt reference patterns embedded either at temporally regular or jittered positions within acoustic sequences (i.e., with a fixed or randomly varying duration of the interval between pattern occurrences) while listeners’ attention was either directed towards or away from the pattern repetitions. As stimulus material we used acoustic patterns that were randomly generated and transformed to match statistical properties of natural sounds (see Methods for details). This allowed us to study learning of initially unfamiliar, complex, and not semantically categorisable sounds that are closer to naturalistic sounds than previously used artificial stimulus material (e.g., white noise) and yet carefully acoustically controlled (other than, e.g., actual environmental sounds). In accordance with previous literature (see previous section), we defined the relevant implicit memory effect as a change in two different EEG markers: Both amplitude of the ERP and EEG phase coherence to repeating pattern onsets within a sequence should differ between the reference pattern that (unbeknownst to the participants) recurred across trials and other patterns that occurred in only one trial throughout the experiment. Note that while we are specifically interested in longer-term pattern learning across multiple trials, the detection of repetitions within trials requires short-term memory representations itself, which likely form the basis for learning across trials. Thus, a particularly critical time window is the first pattern presentation per sequence before any within-trial repetition, since any memory effect at this stage cannot be attributed to short-term, within-trial memory comparisons and is likely related to longer-term memory formation across trials.

We expected that repeated exposure to the reference pattern across multiple trials would lead to the formation of memory representations for this specific acoustic pattern. Such perceptual learning should be reflected in a significant amplitude difference of ERPs to pattern onsets between the learnt reference pattern and other novel, non-recurring patterns, as well as in an increase in ITPC. Moreover, memory formation should be accompanied by an increase of behavioural performance in an auditory pattern repetition detection task. We assumed that both temporal regularity of pattern repetitions and listeners’ attention might shape how acoustic patterns are perceptually learnt. Specifically, it is plausible to hypothesise that the memory effect would be enlarged in temporally regular compared to jittered sequences and when listeners’ attention is direct towards compared to away from the pattern repetitions. Above and beyond a potential modulation by temporal regularity and listeners’ attention, a significant memory effect across conditions would point towards a flexible learning mechanism that enables the formation of robust memories across different, naturally variable learning contexts.

In addition to changes in EEG responses and behavioural performance as our key dependent measures, we assessed active behavioural recognition of learnt patterns as a more direct correlate of memory formation. Participants were asked to complete an unexpected memory test at the end of the experimental sessions in which they had to choose which out of two stimuli they felt they had heard before. This allowed us to test whether the acquired memory representations can become (at least to some degree) actively accessible, and whether more indirect and more direct correlates of learning are similarly or differently influenced by temporal regularity and attention.

## Materials & Methods

### Participants

A total of 29 healthy adults (26 of them female, three male) participated in the study. They were between 18 and 32 years old (*M* = 21.38 years, *SD* = 3.21 years). All of them reported normal hearing, normal or corrected-to-normal vision and no history of any neurological or psychiatric disorder. Three participants were left-handed, the remaining 26 right-handed, as assessed with the short form of the Edinburgh Handedness Inventory (Oldfield, 1971). All participants were psychology undergraduate students and received course credits for their participation. They were naïve regarding the purpose of the study and gave written informed consent before the testing started. All experimental procedures were in accordance with the Declaration of Helsinki and the study was approved by a local ethics committee.

### Stimuli

As auditory stimulus material we used sequences of so-called correlated noise, which is described in detail elsewhere (McDermott et al., 2011). Correlated noise was recently used to study auditory perceptual learning (Ringer et al., 2022a) and comes with the advantage that it is randomly generated, and particular exemplars are unfamiliar to the listeners, while it shares statistical properties with natural sounds. We created correlated noise sequences with a duration of 3500 ms, including 5-ms onset and offset ramps (half-Hanning windows) using the Gaussian Sound Synthesis Toolbox (http://mcdermottlab.mit.edu/Gaussian_Sound_Code_for_Distribution_v1.1) in Matlab (version R2021a; The MathWorks Inc., USA). Randomly generated white noise sequences were transformed using a generative model, such that the resulting correlated noise sequences contained a correlative structure, i.e., correlations of spectral energy values between temporally and spectrally adjacent sampling points in the spectrogram that decreased in strength with increasing temporal and spectral distance. Specifically, the strength of the correlation decreased with -0.065 per time window (20 ms) along the temporal dimension and with -0.075 per frequency window (0.196 octaves) along the spectral dimension. These decay constants were chosen to match our artificially generated stimuli with the correlative structure of natural sounds such as speech stimuli or environmental sounds (McDermott et al., 2011).

In accordance with previous studies using the implicit noise-learning paradigm (e.g., Agus et al., 2010), we created three types of sequences. “Noise” sequences (N) consisted of 3500 ms of random correlated noise, without any repetitions in the sound. In “repeated noise” sequences (RN), a certain 200-ms segment was repeated over the course of the sequence such that it occurred in total six times within the sound. “Reference repeated noise” (RefRN) sequences had the same structure as RN sequences and only differed from them in that they included the exact same repeated pattern in several trials within an experimental block, while all RN and N sequences (and 200-ms patterns contained in them) occurred in just exactly one trial throughout the whole session. 200-ms patterns were created separately for the respective trial (or block in the case of RefRN) and inserted into the 3500-ms sequences at certain time points (see below). As the repetition of a fixed 200-ms pattern throughout RN and RefRN sequences inevitably introduced a local change in the correlative structure of the sounds, we controlled for that by including such local changes also in the N sequences. Concretely, for each N trial, six (different) 200-ms segments were created and inserted into the sequence. That way, the three types of sequences only differed with regard to whether or not a specific pattern occurred repeatedly, but not with regard to local disruptions in the correlative structure. To avoid noticeable clicks at segment boundaries due to abrupt changes in the spectrum, cross-fading (using 5-ms half-Hanning windows centred 2.5 ms relative to the beginning and -2.5 ms relative to the end of an inserted 200-ms segment) was applied at all transitions in all sequences. In addition to the manipulation of the within- and across-sequence pattern repetitions (N, RN, RefRN), sequences varied with regard to the temporal Regularity of the repetitions of the repeated 200-ms pattern, such that pattern onsets within the sequence were either regular or jittered. In the regular sequences, patterns were repeated with a constant interval of 300 ms between patterns. In the jittered sequences, patterns were repeated at variable intervals, such that each interval between two consecutive pattern occurrences was randomly chosen between 50 ms and 550 ms (around a mean of 300 ms), with the restriction that two adjacent inter-pattern intervals must differ in duration by at least 50 ms. Across both regular and jittered sequences, the first pattern started at a randomly selected time point between 50 ms and 500 ms relative to sound onset. The structure of the acoustic sequences in the different conditions is illustrated in Figure 1A. Audio files with example stimuli can be found in the online supplemental material (https://osf.io/r8gea/?view_only=4580e185befa474eac7a2d0efb3c615e).

**Figure 1.**
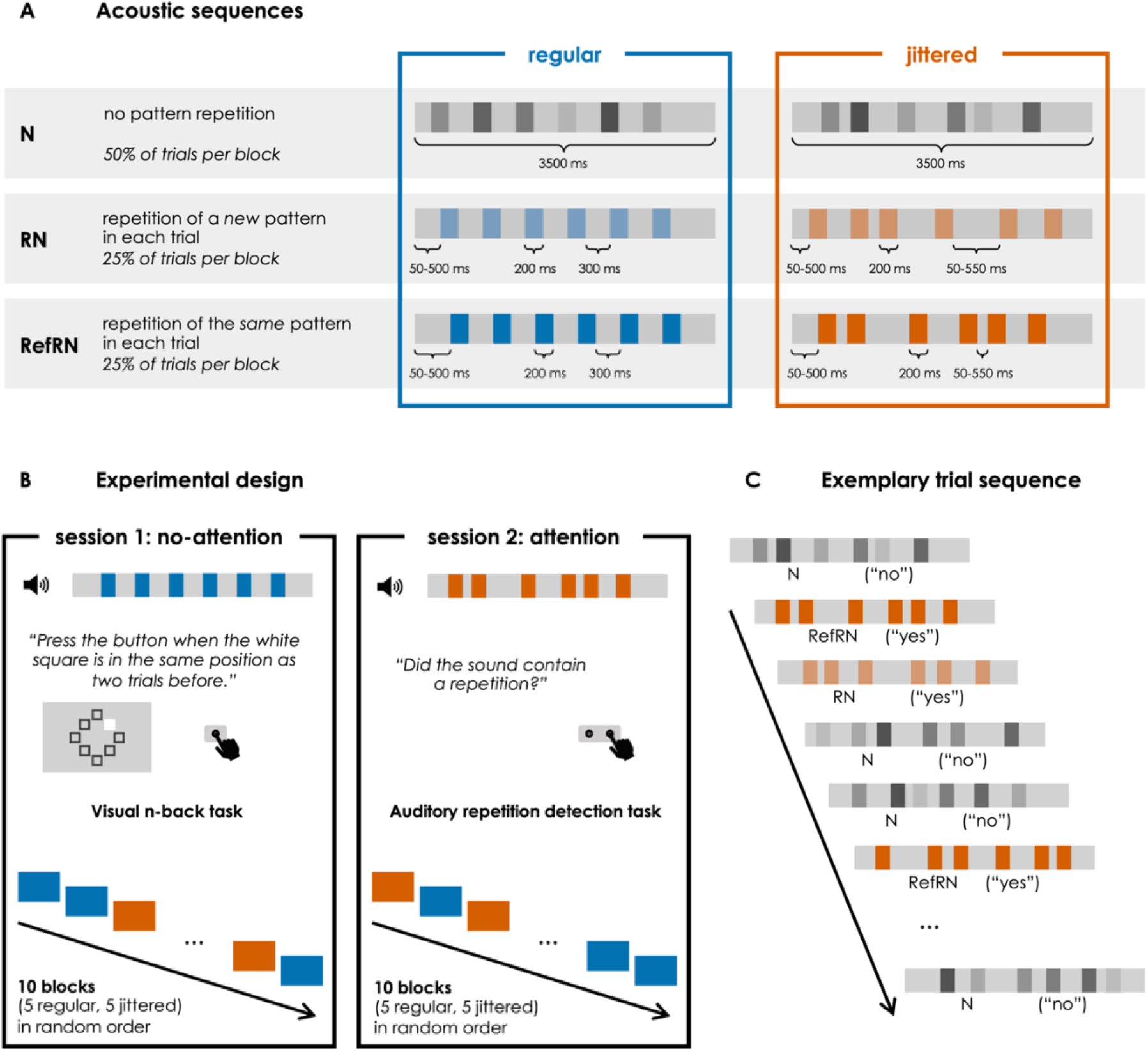
A: Illustration of the acoustic stimulus sequences. B: Experimental design. Participants took part in two EEG sessions (in a fixed session order). Different reference patterns were used in both sessions. C: Exemplary trial sequence. Trials were presented in a random order, with the only restriction that two RefRN trials must not immediately follow each other.

### Procedure

#### General procedure

Each participant completed two EEG sessions, each of which lasted between two and three hours. The two sessions took place on separate days, with an average interval of 13 days in between. Attention was manipulated between sessions, such that in one session the listeners’ attention was directed towards and in the other session away from the pattern repetitions within the auditory sequences. In the first session (no-attention), they performed a visual distractor task while instructed to ignore the auditory sequences. In the second session (attention), they were asked to perform a repetition detection task and indicate after each sequence whether or not it contained a repetition. This session order was fixed for all participants in order to avoid that they were aware of the repetitions in the auditory stimulation in the no-attention session after performing the auditory repetition detection task in the session before. As previous studies suggested that memory representations of learnt patterns persist over several weeks (Agus et al., 2010; Bianco et al., 2020; Viswanathan et al., 2016), different patterns were used for RefRN sequences in both sessions (while the same RN and N sequences, which occurred only once per session, were used for both sessions). The experimental design and an exemplary trial sequence are illustrated in Figure 1B and 1C.

During the experiment, participants were seated in an acoustically and electrically shielded chamber. Instructions for the respective task and visual stimuli were displayed on a computer screen behind a window outside the cabin at approximately 80 cm distance from the participants’ eyes. Auditory stimuli were delivered binaurally via headphones (Sennheiser HD-25-1, Sennheiser GmbH & Co. KG, Germany). Stimulus presentation and response registration was controlled using the Psychophysics Toolbox extension (PTB-3; Brainard, 1997; Kleiner et al., 2007) in GNU Octave (version 5.2.0; Eaton et al., 2020). Participants’ behavioural responses were captured with a response time box (Suzhou Litong Electronic Co., China).

In each session, participants completed ten experimental blocks, five with regular and five with jittered sequences, in a random order. One block lasted about six minutes and breaks could be taken between blocks as required. Within each block, 60 auditory sequences were presented, 50% of which were N sequences, and 25% RN and RefRN sequences, respectively. Sequence order was pseudorandomised with the restriction that two RefRN sequences must not immediately follow each other. The duration of the silent interval between two sequences was jittered between 2175 and 2625 ms (in steps of 50 ms) around a mean of 2400 ms.

#### Repetition detection task in the attention session

In the second session (attention), participants were asked to perform a repetition detection task, deciding for every sequence whether it contained a repetition or not. At the beginning of the session, they had the chance to familiarise themselves with the different types of sequences, which were introduced to them as having either “regular” repetitions (RN/RefRN regular), “irregular” repetitions (RN/RefRN jittered), or no repetition at all (N regular/jittered). An example sequence (which was not part of the actual stimulus pool) was provided for each of the three types of sequences and participants could listen to the examples as often as they wanted. They were informed that some blocks would contain only regular and other blocks only irregular sequences, and that 50 % of the sequences per block contained repetitions. During auditory sequence presentation, a white fixation cross was displayed in the centre of the screen against a grey background. After the end of the sequence, the two response options (repetition/no repetition) were displayed on the screen until a response was given or a maximum response interval of 2000 ms expired. Participants gave their responses by pressing either the left or the right button on the response time box (with the corresponding index finger). Response sides were counterbalanced across participants. At the end of each block, feedback, i.e., the percentage of correct responses, was provided.

#### Visual distractor task in the no-attention session

In the first session (no-attention), additional visual stimulation was delivered concurrently. The visual display consisted of a white fixation cross against a grey background in the centre of the screen, surrounded by eight squared frames (width/height: 0.50° visual angle) that were arranged in a circle (radius: 2.11° visual angle) at equal distance from each other. In each block, 240 visual trials were presented with no temporal or other systematic relationship to the auditory stimulation. In each visual trial, a white square appeared for 150 ms at one of the eight frame positions.

Participants were instructed to monitor the appearances of the white square, while fixating the cross in the centre, and press a button as quickly as possible whenever the current position of the white square was identical to the position two trials before (2-back task). The visual stimulus onset asynchrony was jittered between 1425 and 1575 ms (in steps of 10 ms) around a mean of 1500 ms. 2-back targets occurred randomly in 10 % of the trials, with the restrictions that the first five trials per block were always non-targets and each target was followed by at least two non-targets. Targets occurred equally often at each of the eight frame positions, while the square positions were randomly chosen for the non-target trials. To engage the participants in the visual distractor task at the beginning of the block, auditory stimulation began only five seconds after the visual stimulation. Before the actual experiment started, participants completed a short training (24 trials) on the visual task in the absence of concurrent auditory stimulation. During this training, feedback was provided in each trial (correct/incorrect; displayed below the circle of squared frames for 500 ms), while during the ten experimental blocks feedback (percentage of hits and false alarms, mean reaction time) was only provided at the end of each block.

#### Active auditory memory test

At the end of each session, participants were asked to complete an additional unexpected two-alternative forced-choice auditory memory test. In each trial of this test block, they were presented with two 200-ms patterns, separated by 1000 ms of silence, one of which was a previously learnt reference pattern and the other one a random unfamiliar pattern. They should then select the pattern that they felt they had already heard before during one of the ten blocks. As in the repetition detection task, the silent inter-trial interval (between the offset of the second sound and the onset of the first sound of the following trial) was jittered between 2175 and 2625 ms and the maximum response interval was 2000 ms. The order of the reference pattern (i.e., target) and the random filler stimulus (i.e., non-target) was counterbalanced across trials, such that the reference noise did not occur in the same position in more than three trials in a row. Each of the ten reference patterns was presented four times during the test block. Patterns were presented in the same order as during the experimental blocks before and this fixed order (RefRN1, RefRN2, RefRN3, …) was repeated four times, in order to avoid that the same reference pattern could occur in two immediately successive trials. To make sure that participants could not base their responses solely on pattern repetition within the test block, we used ten random filler patterns as non-targets and presented each of them equally often as each reference pattern (i.e., four times). These filler patterns were assigned to the trials pseudo-randomly, such that the same stimulus did not occur in two trials in a row. No feedback was provided in the test block.

### EEG data acquisition

EEG was recorded continuously from 64 active Ag/AgCl electrodes mounted in an elastic cap according to the extended international 10-20 system. Four electrodes were placed on the outer canthus of each eye and above and below the right eye to capture horizontal and vertical eye movements. Additionally, signals were recorded from left and right mastoids (M1, M2) and one electrode placed on the tip of the nose served for later offline referencing. During preparation, we made sure that all electrode offsets were kept below 30 μV. Signals (referenced to the CMS-DRL ground) were amplified with a BioSemi ActiveTwo amplifier (BioSemi B.V., Amsterdam, The Netherlands) and digitised with a sampling rate of 512 Hz.

### Data analysis and statistical inference

#### Behavioural data

Analysis of behavioural data was done in RStudio (version 4.0.2, RStudio Inc., USA).

### Visual distractor task

We computed hit rates (proportion of correctly detected 2-back targets), false alarm rates (proportion of non-targets erroneously classified as targets) and mean reaction times (of correct target detections) across blocks for each participant. Means and standard deviations across individuals are reported for these three performance measures to demonstrate that, on average, participants complied with the task instructions and performed above chance in the visual task.

### Repetition detection task

Repetition detection performance was quantified within the framework of signal detection theory, an approach that is commonly used to separate response accuracy from response bias in perceptual categorisation tasks (Macmillan, 2001). Hits were defined as the trials in which the sequence contained repetitions and participants correctly responded that they heard a repetition, and false alarms as trials in which the sequence did not contain repetitions, but participants erroneously responded that they heard a repetition. For each participant, we computed d’ sensitivity indexes for RefRN and RN sequences from the respective hit rates and the false alarm rates in regular and jittered blocks, respectively. To avoid computation with infinite values, we applied the so-called log-linear transformation (Hautus & Lee, 2006).

A two-way repeated-measures ANOVA (implemented in the package “ez”; Lawrence, 2016) with the factors Repetition Type (RefRN, RN) and Regularity (regular, jittered) was used to statistically test for a memory effect and its modulation by temporal regularity. Substantial learning of reference noises, i.e., a significantly higher sensitivity for RefRN compared to RN, would be reflected in a significant main effect of Repetition Type, and a substantial modulation by temporal regularity in a significant interaction with Regularity. Greenhouse-Geisser correction was applied to correct for non-sphericity (as indicated by a significant Mauchly’s test with *p* < .05). Statistical significance was defined by the standard .05 alpha criterion. In addition to the frequentist tests, we computed complementary Bayesian tests and report Bayes Factors (BF_10_), using the package “BayesFactor” (Morey & Rouder, 2011; Morey et al., 2018; Rouder et al., 2009, 2012). Reported Bayes Factors were computed by comparing evidence for models that include the effect of interest against reduced matched models that do not include the effect of interest (in accordance with recommendations by Bergh et al., 2020). In accordance with widely used conventions (Lee & Wagenmakers, 2014), BF_10_ > 3 (10) was considered moderate (strong) evidence for the alternative hypothesis and BF_10_ < 0.33 (0.1) was considered moderate (strong) evidence for the null hypothesis, while values in between were deemed inconclusive.

### Memory test

Active pattern recognition performance was measured as the percentage of correct responses in the memory test. First, performance was tested against 50-% chance level using a one-sided one-sample *t*-test for the regular and jittered condition in the attention and no-attention session, respectively. Second, we compared the performance between conditions by means of a two-way repeated-measures ANOVA with the factors Regularity (regular, jittered) and Attention (attention, no-attention). Where applicable, a correction for non-sphericity was used as described above, and the frequentist analysis was again complemented with a Bayesian ANOVA.

#### EEG data

EEG data were processed offline in Matlab (version R2022a), using the EEGLAB (version 14.1.2; Delorme & Makeig, 2004) and the FieldTrip (Oostenveld et al., 2011) toolbox. Subsequent statistical analysis of mean amplitudes was done in RStudio (version 4.0.2, RStudio Inc., USA).

### Pre-processing

Data were pre-processed separately for each EEG recording session (i.e., two per participant). First, data were referenced to the channel located on the tip of the nose. Noisy channels were excluded from pre-processing and later spherically spline interpolated (as a last pre- processing step) if their signal variance exceeded an absolute z-score of 3.0. Data of the remaining channels were high-pass and low-pass filtered with Kaiser-windowed sinc finite impulse response (FIR) filters at 0.2 Hz (transition bandwidth: 0.4 Hz, maximum passband deviation: 0.001, filter order: 4638) and 35 Hz (transition bandwidth: 5 Hz, maximum passband deviation: 0.001, filter order: 372).

The filtered continuous data were cut into epochs that ranged from -100 ms to 4000 ms relative to the onset of the 3500-ms auditory sequences. We then used an independent component analysis (ICA) to clean the epoched data from physiological and technical artefacts. To improve signal-to-noise ratio for decomposition, ICA was computed on a copy of the data filtered with a 1-Hz high-pass filter (transition bandwidth: 0.5 Hz, maximum passband deviation: 0.001, filter order: 3710) and the same 35-Hz lowpass filter. Data were epoched as described above and epochs with a peak-to-peak difference that exceeded 750 µV were discarded. To shorten computation time, data were down- sampled to 128 Hz before the ICA decomposition. ICA weights were obtained with an infomax algorithm implemented in EEGLAB’s runica function and transferred to the dataset that was pre- processed using the final filter parameters. Independent components were classified using the IC Label plugin for EEGLAB (Pion-Tonachini et al., 2019) and artefactual components classified as eye blinks, muscle activity, cardiac activity, line noise or channel noise were automatically selected and removed. To minimise influences of visual processing and motor responses on auditory event-related potentials during the visual distractor task in the first session (no-attention), auditory events within 500 ms before and after a button press as well as within the 500-ms interval prior to visual targets were excluded from the analysis.

### Event-related potential analysis

For an overview of the electrophysiological response to the full auditory sequence, we extracted epochs that ranged from -100 ms to 3000 ms relative to the onset of the first 200-ms pattern, covering all six pattern presentations per sequence. Epochs were baseline-corrected to the 100-ms interval prior to the first pattern onset. Any epoch with a peak-to-peak difference that exceeded 300 µV was excluded from the analysis. The remaining epochs were re-referenced to the algebraic mean of both mastoid electrodes (M1, M2) and, for each individual, averaged for N, RN and RefRN sequences in each Regularity and Attention condition, respectively. From these within-participant averages, grand averages were computed for each condition (using the grandaverage plugin for EEGLAB, authored by Andreas Widmann; https://github.com/widmann/grandaverage).

For the analysis of pattern-related responses to single pattern occurrences, we extracted epochs that ranged from -100 ms to 500 ms relative to the onset of each 200-ms pattern. Epochs were baseline-corrected to the 100-ms interval prior to pattern onset. Any epoch with a peak-to-peak difference that exceeded 150 µV was excluded from the analysis, and the remaining epochs were re-referenced to the algebraic mean of both mastoid electrodes. Computation of within-participant averages and grand averages followed the same procedure as described above for the response to the full sequence. Subsequent analysis steps were done separately for different pattern positions within the sequence: a) averaged across position two to six, i.e., the first to the fifth pattern repetition per sequence, and b) for the first presentation per sequence only, i.e., before any within-sequence pattern repetition.

Time windows of interest for the statistical comparison of mean amplitudes between conditions were defined using a non-parametric cluster-based permutation approach. Specifically, we performed a cluster-based permutation test to identify clusters of significant amplitude differences between RefRN and RN at temporally and spatially adjacent samples (Maris, 2012; Maris & Oostenveld, 2007). In accordance with recent recommendations on how to avoid bogus effects due to selective choice of analysis parameters in ERP research (Luck & Gaspelin, 2017), we collapsed RefRN and RN waveforms across the four Regularity and Attention conditions and fed these condition averages into the cluster-based permutation test. That way, we reduce the risk that potential differences between Regularity and Attention conditions are merely a result of a biased selection of the time window for statistical analysis. The cluster-based permutation test was performed in a time window ranging from 0 to 500 ms relative to pattern onset, applying an alpha level and cluster alpha of 0.05 and using a Monte Carlo approximation with 1000 permutations to estimate cluster-level significance probability. Based on the results of the cluster-based permutation tests, we identified relevant time windows for analysis in which a memory effect occurs (on average across conditions) that ranged from 140 to 220 ms relative to pattern onset for the average across pattern position two to six, and from 260 to 500 ms for the first pattern presentation per sequence.

From these time windows, mean RefRN and RN amplitudes were extracted for each of the four Regularity and Attention conditions at electrode Fz (since both the current data and previous studies suggested a fronto-central distribution of the memory effect in the ERP, e.g., Andrillon et al., 2015; Hodapp & Grimm, 2021). Moreover, RefRN-minus-RN difference waveforms were computed for the four Regularity and Attention conditions.

Statistical evaluation of mean amplitudes was done independently for the average across pattern position two to six and the first pattern presentation per sequence. We used a three-way repeated-measures ANOVA with the factors Repetition Type (RefRN, RN), Regularity (regular, jittered) and Attention (attention, no-attention), respectively. A significant main effect of Repetition Type would indicate a substantial memory effect, and any significant interaction with Repetition Type a substantial modulation of the memory effect by the respective factor. Where applicable, a correction for non-sphericity was used as described above, and we again computed both frequentist and Bayesian tests.

### Time-frequency analysis

For the time-frequency analysis, we extracted epochs that ranged from -200 ms to 800 ms relative to within-sequence pattern repetition onsets, i.e., the onsets of the 200-ms pattern at position two to six. Epochs were de-meaned and any epoch with a peak-to-peak difference higher than 150 µV was excluded from the analysis. We applied a convolution with Morlet wavelets to extract phase information from single epochs, with 1500 ms of zero-padding at both ends of the epoch. Parameters of the Morlet wavelet were adjusted linearly from three to seven wavelet cycles over a frequency range from 1 to 10 Hz in steps of 0.2 Hz. From the results of the wavelet convolution, we computed ITPC (between individual epochs) at electrode Fz for each participant at each frequency and sampling point, separately for RefRN and RN in each Regularity and Attention condition. For statistical analysis, a frequency window of interest ranging from 3 to 6 Hz was chosen based on previous literature (e.g., Luo et al., 2013) and visual inspection of the spectrograms of the current data. Note that this low-frequency range does on purpose not include 2 Hz as the stimulation frequency in the regular condition, in order to avoid a bias between Regularity conditions that is driven by direct entrainment to the stimulus rhythm in regular sequences. Analogous to the ERP analysis, the time window of interest was defined using a cluster-based permutation test performed on RefRN and RN waveforms collapsed across the four Regularity and Attention conditions between 0 and 500 ms relative to pattern onset (see above for detailed parameters of the cluster-based permutation test). Based on the results of the test, we identified a relevant time window of RefRN-minus-RN differences that ranged from 0 to 190 ms relative to pattern onset.

From this time window, we extracted mean ITPC in the frequency range from 3 to 6 Hz for RefRN and RN in each of the four Regularity and Attention conditions. ITPC coefficients were statistically compared by means of a three-way repeated-measures ANOVA with the factors Repetition Type (RefRN, RN), Regularity (regular, jittered) and Attention (attention, no-attention). Where applicable, a correction for non-sphericity was used as described above, and we again computed both frequentist and Bayesian tests.

## Results

### Behavioural data

#### Visual distractor task

Participants detected on average 74.3 ± 11.8 % (*M* ± *SD*) of the visual 2-back targets. False alarms occurred in 2.4 ± 2.8 % of the non-target trials. The mean reaction times in the trials in which participants correctly detected n-back targets was 608 ± 66 ms. These data suggest that the task was challenging, and performance was rather far from perfect, but participants, on average, performed well above chance. Thus, we conclude that they complied with the instruction to focus on the visual task and the task successfully directed their attention towards the visual (and away from the auditory) stimulation.

#### Repetition detection task

Repetition detection performance is shown in Figure 2. On average, listeners detected pattern repetitions above chance in both RN and RefRN sequences across regular and jittered blocks, with no significant overall performance difference between regular and jittered repetitions (main effect of Regularity: *F*(1, 28) = 3.22, *p* = .084, partial η^2^ = .10, BF_10_ = 2.15). Importantly, we found a significant memory effect across regularity conditions (main effect of Repetition Type: *F*(1, 28) = 29.64, *p* < .001, partial η^2^ = .51, BF_10_ = 224.67). Thus, recurrence of reference patterns across multiple trials led to an increase in repetition detection performance (i.e., a higher sensitivity index d’) for RefRN sequences relative to RN sequences, which contained patterns that only occurred in exactly one trial. This memory-related performance increase was not substantially modulated by the temporal regularity of pattern repetitions within a sequence (Repetition Type x Regularity interaction: *F*(1, 28) = 1.99, *p* = .169, partial η^2^ = .07, BF_10_ = 0.46).

**Figure 2.**
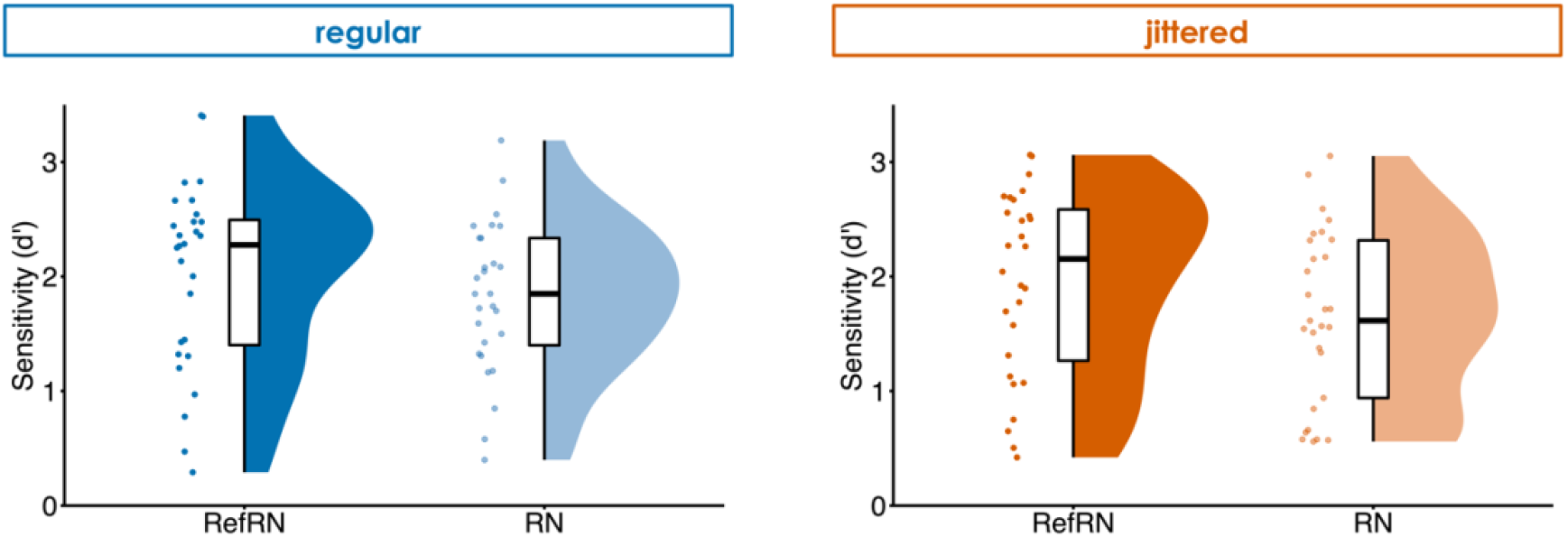
Behavioural repetition detection performance in the attention session, measured by the sensitivity index d’, separately for RefRN (containing repetitions of the reference pattern that reoccurred across trials) and RN sequences (containing repetitions of a pattern that occurred in only one trial throughout the experiment) in regular and jittered blocks. Error bars indicate ± 1 standard error of means (SEM).

#### Active memory test

Performance in the active memory test did not significantly exceed 50 % in either Regularity or Attention condition (all *p*’s > .196, all BF_10_’s < 0.45). Thus, participants were unable to recognise previously learnt reference patterns above chance. Moreover, recognition performance was unaffected by manipulations of the temporal regularity of pattern repetitions during learning and listeners’ attention towards the pattern repetitions (both main effects and Regularity x Attention interaction: all p’s > .134, all BF_10_’s < 0.82).

### EEG data

#### Event-related response to the full sequence

As shown in Figure 3, a negative sustained potential emerged in all conditions after the onset of the first pattern. Note that the absence of a typical sound onset response in the average ERP is due to the variable interval between sound onset and first pattern onset (50 to 500 ms) across trials. Across conditions, amplitudes of the sustained potential were larger in the attention compared to the no-attention session, in line with earlier reports of increased sustained potential amplitudes in response to attended compared to ignored auditory stimuli (Picton et al., 1978). On top of the sustained response, the isochronous onsets of the repeating pattern (i.e., every 500 ms) in RN and RefRN sequences were reflected in a periodic amplitude modulation in regular blocks independent of the listeners’ attention. In jittered blocks, any response time-locked to the onset of the repeating pattern is levelled out on average across trials due to phase shifts between sequences. A difference between RefRN and RN sequences, i.e., a memory effect through recurrence of the reference pattern across trials, is visible in the attention session (across both regular and jittered blocks), most pronounced between around 250 and 1500 ms relative to the first pattern onset. No such memory-related amplitude difference between RefRN and RN sequences is observable in the no-attention session.

**Figure 3.**
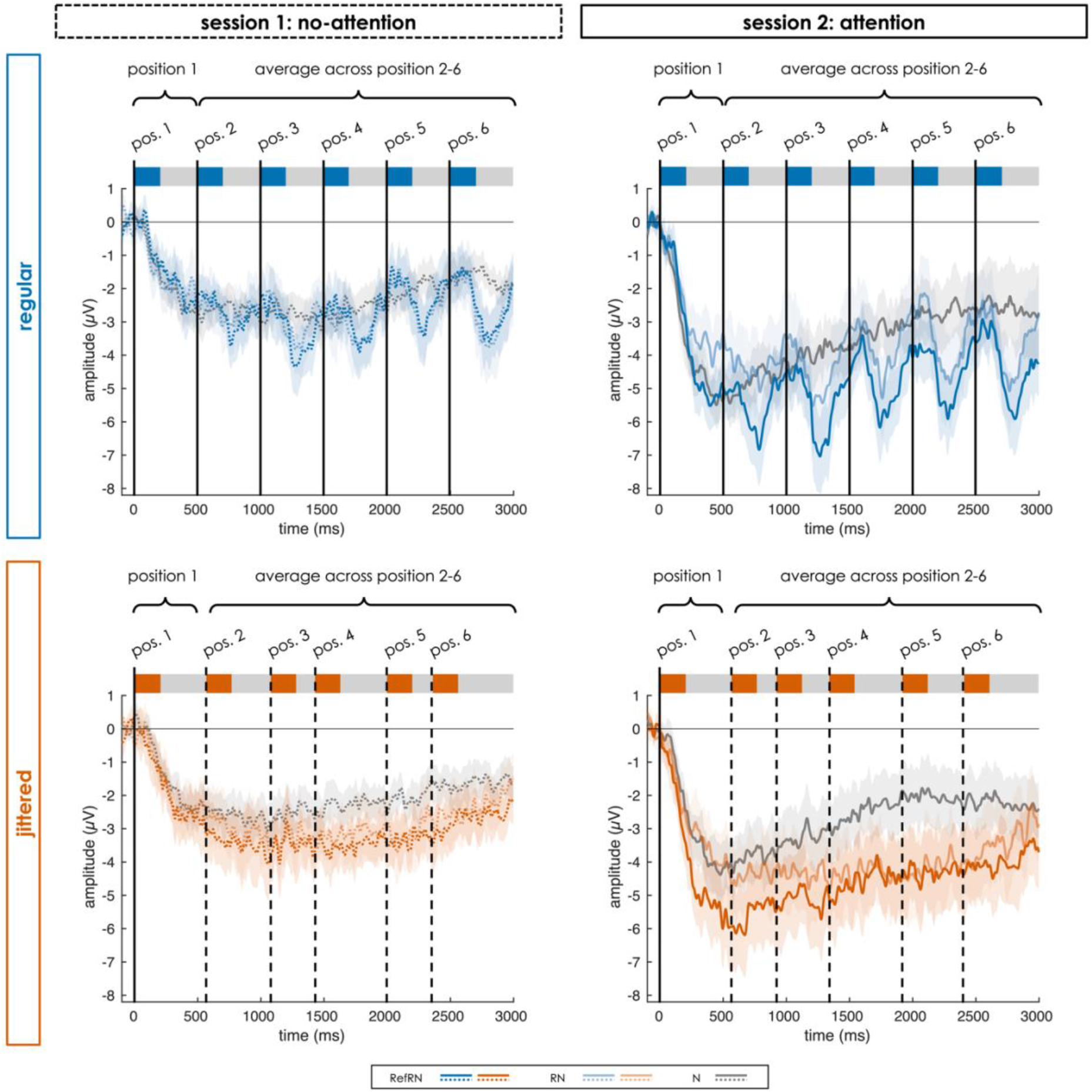
ERP response to the full auditory sequence relative to the onset of the first pattern per sequence (0 ms) at electrode Fz for N, RN and RefRN sequences in the four Regularity (regular, jittered) and Attention (attention, no-attention) conditions. The structure of the stimulus sequences is illustrated schematically above the ERP plot, with vertical lines indicating pattern onset and blue/orange boxes representing the repeating pattern (in RN and RefRN sequences) within the continuous sequence (grey box). Note that the pattern onset times in the jittered condition only refer to one exemplary trial and were actually jittered between individual sequences. Shaded areas indicate ± 1 SEM.

#### Event-related responses to pattern repetitions at position 2-6

As shown in Figure 4A, repetitions of a certain 200-ms pattern in RN and RefRN sequences elicited a pattern-related response that was time-locked to the onset of the repeating pattern at position two to six within the sequence (i.e., the first to the fifth pattern repetition). In all Regularity and Attention conditions, the pattern-related response was characterised by a prominent negativity between around 150 and 400 ms, with a peak between 250 and 300 ms, relative to pattern onset, preceded by a weaker positivity within the first 150 ms after pattern onset. No such characteristic modulation of the ERP was observed for N sequences, suggesting that the response is in fact related to the repetition of a specific pattern and not (only) to the disruption of the local correlative structure of the acoustic sequence. Especially in the attention session, the negativity tended to have a narrower peak shape and peak amplitudes of the negativity tended to be higher for RefRN compared to RN sequences, resulting in a somewhat steeper slope.

**Figure 4.**
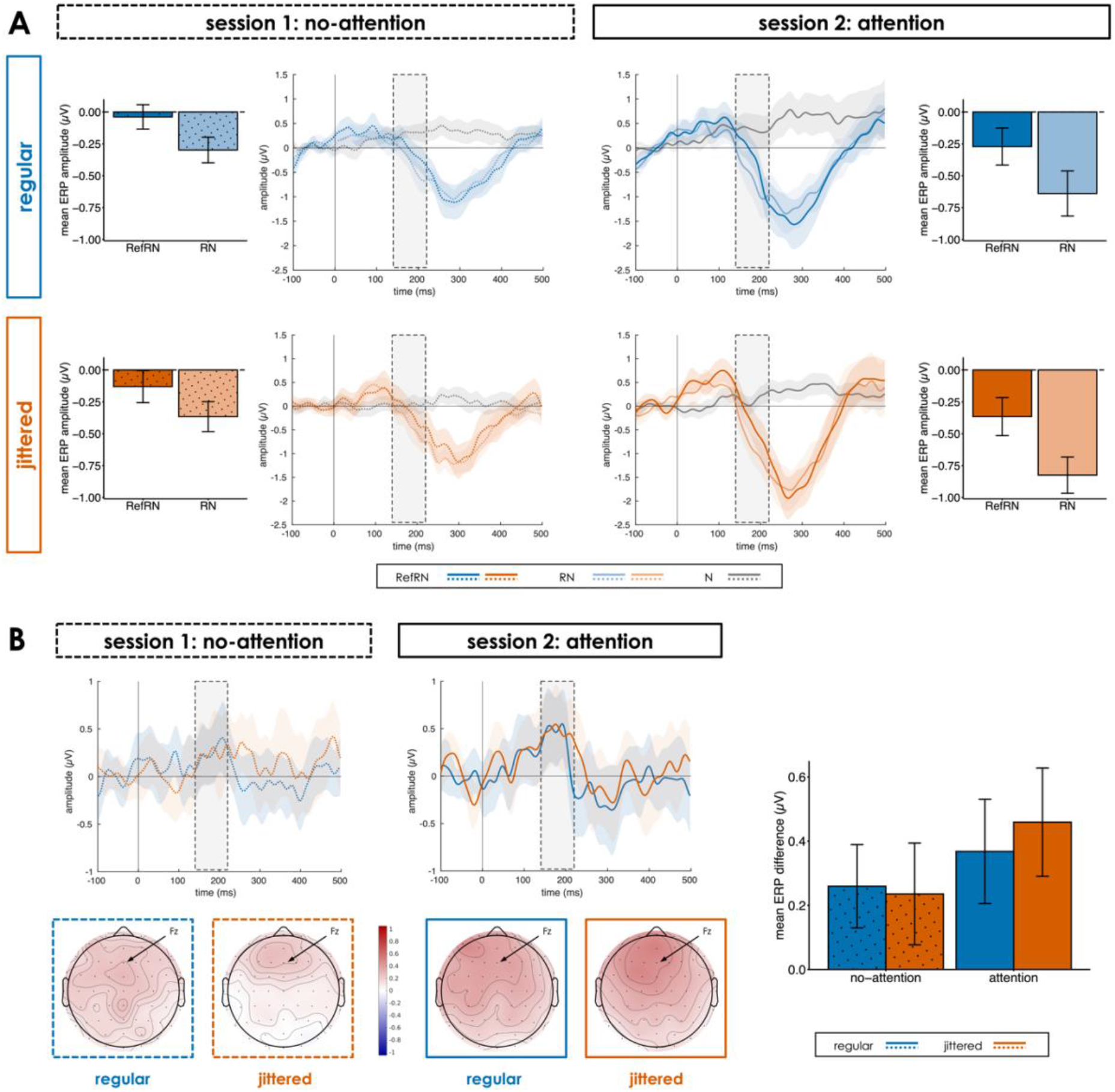
A: Middle panels: Pattern-related response relative to the onset of the repeating pattern at position two to six within the sequence (0 ms) at electrode Fz for N, RN and RefRN sequences in the four Regularity (regular, jittered) and Attention (attention, no-attention) conditions. Outer panels: Mean amplitudes in the time window of interest (140 to 220 ms relative to the first pattern onset; marked with dotted boxes) for RefRN and RN sequences. B: Upper row: Difference waveforms (RefRN-minus-RN) for the four Regularity and Attention conditions. Lower row: Topographies of the RefRN-minus-RN difference in the time window of interest. Right panel: Mean amplitudes of the difference waveforms. Shaded areas in the ERP plots and error bars in the bar plots indicate ± 1 SEM.

#### Pattern-related ERP response

In the previously identified time window of interest between 140 and 220 ms after pattern onset, we found a significant difference in amplitude between RefRN and RN sequences (main effect of Repetition Type: *F*(1, 28) = 15.80, *p* < .001, partial η^2^ = .36, BF_10_ = 199.01). Amplitudes during this time window in the slope between positive and negative peak were less negative for RefRN compared to RN sequences, likely related to the narrower peak shape. As illustrated in Figure 4B, the memory effect consistently pointed to this direction and exhibited a fronto-central topography across all four Regularity and Attention conditions. The ANOVA results suggested that the memory effect was not substantially modulated by Regularity (Repetition Type x Regularity interaction: *F*(1, 28) = 0.04, *p* = .848, partial η^2^ < .01, BF_10_ = 0.22), Attention (Repetition Type x Attention interaction: *F*(1, 28) = 1.22, *p* = .279, partial η^2^ = .04, BF_10_ = 0.32) or an interaction between both factors (Repetition Type x Regularity x Attention interaction: *F*(1, 28) = 0.20, *p* = .656, partial η^2^ = .01, BF_10_ = 0.23). Beyond the memory effect of interest, amplitudes were overall larger in the attention compared to the no-attention session (main effect of Attention: *F*(1, 28) = 6.44, *p* = .017, partial η^2^ = .19, BF_10_ = 112.62), but not significantly influenced by Regularity (main effect of Regularity: *F*(1, 28) = 1.92, *p* = .177, partial η^2^ = .06, BF_10_ = 0.33; Regularity x Attention interaction: *F*(1, 28) = 0.25, *p* = .623, partial η^2^ = .01, BF_10_ = 0.23).

#### ITPC

As depicted in Figure 5, ITPC was overall highest at low frequencies around 2 Hz within a time range around pattern onset. Crucially, we found differences in ITPC between RefRN and RN sequences in a frequency window beyond the stimulation frequency (i.e., 2 Hz), covering low frequencies between 3 and 6 Hz, which can be attributed to learning of reference patterns.

**Figure 5.**
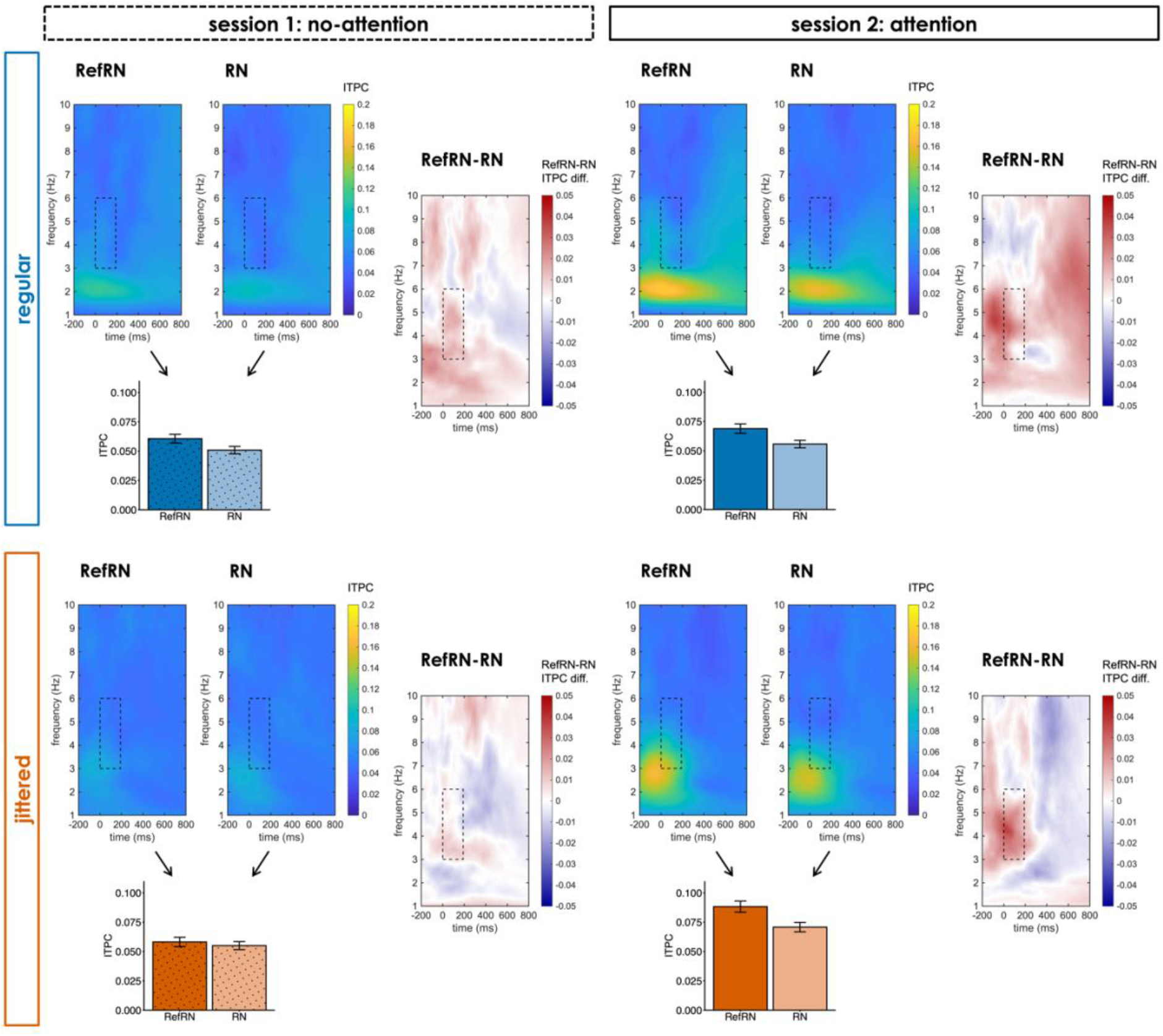
Results of the phase coherence analysis at electrode Fz for the four Regularity (regular, jittered) and Attention (attention, no-attention) conditions. For each condition: Left side, upper panels: Inter-trial phase coherence (ITPC) over frequencies and time relative to pattern onset (0 ms) for RefRN and RN sequences. Left side, lower panel: Mean ITPC extracted from the time-frequency window of interest (3 to 6 Hz, 0 to 190 ms relative to pattern onset; marked with dotted boxes) for RefRN and RN sequences. Right side: RefRN-minus-RN differences in ITPC over frequencies and time. Error bars in the bar plots indicate ± 1 SEM.

In the time window of interest ranging from 0 to 190 ms relative to pattern onset, ITPC was stronger in RefRN compared to RN sequences (main effect of Repetition Type: *F*(1, 28) = 11.52, *p* = .002, partial η^2^ = .29, BF_10_ = 824.96). This memory effect might have been influenced by Attention, such that it was stronger in the attention than in the no-attention session, although Bayesian evidence for a modulation by Attention was inconclusive (Repetition Type x Attention interaction: *F*(1, 28) = 5.30, *p* = .029, partial η^2^ = .16, BF_10_ = 0.93. In contrast, Regularity did not substantially modulate the memory effect (Repetition Type x Regularity interaction: *F*(1, 28) = 0.06, *p* = .801, partial η^2^ < .01, BF_10_ = 0.40; Repetition Type x Regularity x Attention interaction: *F*(1, 28) = 1.33, *p* = .259, partial η^2^ = .05, BF_10_ = 0.17). Beyond the memory effect, phase coherence was overall increased in the attention compared to the no-attention session (main effect of Attention: *F*(1, 28) = 21.90, *p* < .001, partial η^2^ = .44, BF_10_ = 3.79*10^5^). Moreover, ITPC was stronger in jittered than in regular blocks (main effect of Regularity: *F*(1, 28) = 18.13, *p* < .001, partial η^2^ = .39, BF_10_ = 42.95), especially so in the attention session (Regularity x Attention interaction: *F*(1, 28) = 11.47, *p* = .002, partial η^2^ = .29, BF_10_ = 27.75). This incidental finding might be related to the structure of the jittered sequences that (with maximal and minimal intervals between pattern onsets of 750 and 250 ms, respectively) could activate a broader range of frequencies (1.33 to 4 Hz), occasionally (by chance) in a coherent manner across several trials. Thus, other than for regular sequences, the frequency window of interest for the ITPC analysis (spanning 3 to 6 Hz) partly overlaps with possible local stimulation frequencies in jittered sequences.

### Event-related response to the first pattern presentation at position 1

At the first pattern position within the sequence, pattern onset was followed by a negative deflection that emerged around 100 ms after pattern onset in all conditions (see Figure 6A). The negative shift in potential across conditions is due to the first pattern presentation falling into an early phase of the sequence during which the sustained response is building up before reaching its sustained phase. Critically, we found condition differences between RefRN and RN on top of this overall negative shift even before any within-sequence pattern repetition, thus the effect likely reflects pattern recognition based on across-trial memories.

**Figure 6.**
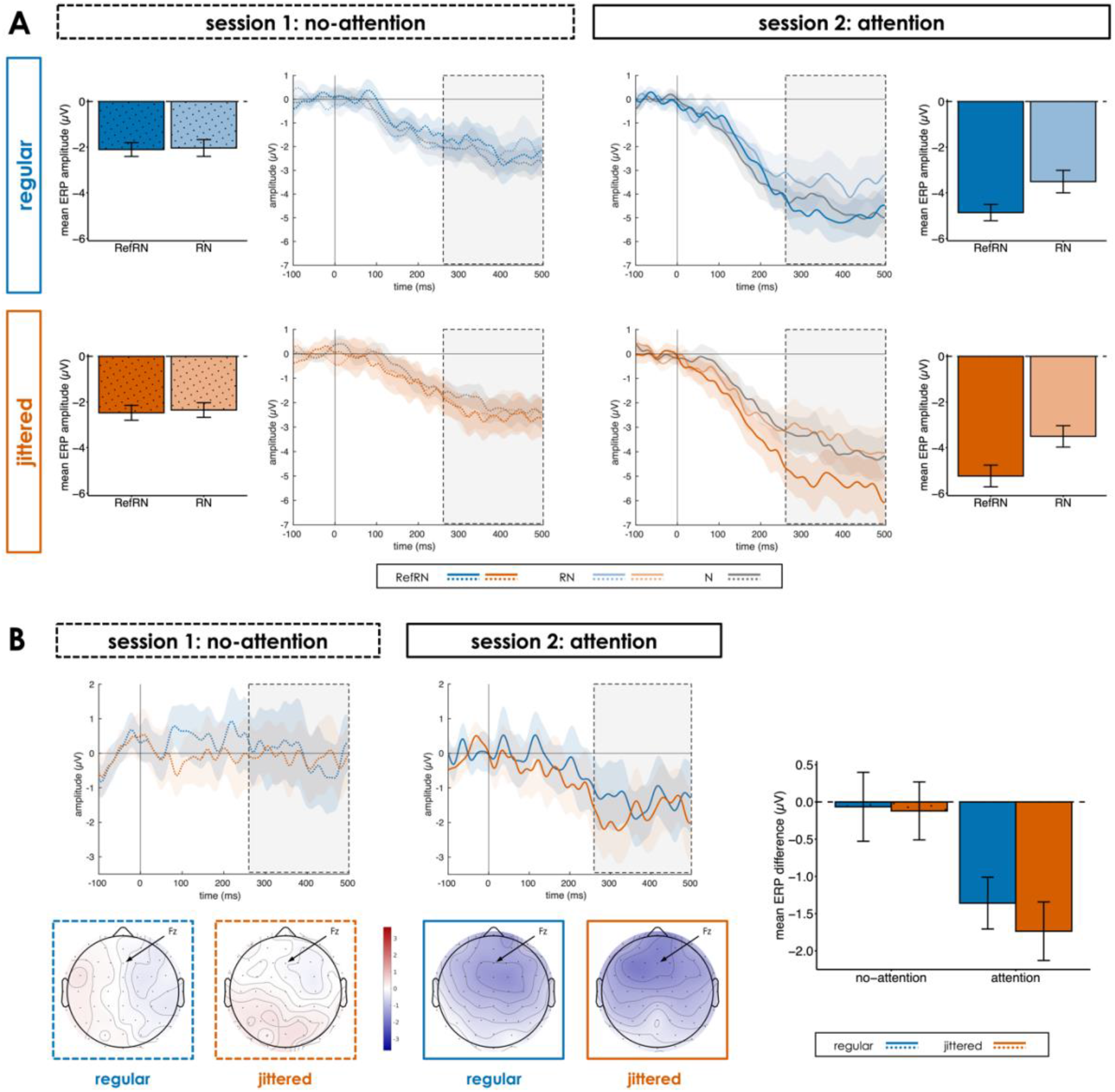
A: Middle panels: Pattern-related response relative to pattern onset at the first pattern position within the sequence (0 ms) at electrode Fz for N, RN and RefRN sequences in the four Regularity (regular, jittered) and Attention (attention, no-attention) conditions. Outer panels: Mean amplitudes in the time window of interest (260 to 500 ms relative to the first pattern onset; marked with dotted boxes) for RefRN and RN sequences. B: Upper row: Difference waveforms (RefRN-minus- RN) for the four Regularity and Attention conditions. Lower row: Topographies of the RefRN-minus- RN difference in the time window of interest. Right panel: Mean amplitudes of the difference waveforms. Shaded areas in the ERP plots and error bars in the bar plots indicate ± 1 SEM.

In the time window of interest between 260 and 500 ms after pattern onset, we found a significant amplitude difference between RefRN and RN sequences (main effect of Repetition Type: *F*(1, 28) = 13.27, *p* = .001, partial η^2^ = .32, BF_10_ = 81.00). This memory effect was substantially modulated by Attention (Repetition Type x Attention interaction: *F*(1, 28) = 16.36, *p* < .001, partial η^2^ = .37, BF_10_ = 34.29). Specifically, only in the attention session amplitudes of the negative potential were larger for RefRN compared to RN sequences, and this effect was again fronto-centrally distributed (as illustrated in Figure 6B). Beyond that, the memory effect was not further modulated by Regularity (Repetition Type x Regularity interaction: *F*(1, 28) = 0.35, *p* = .560, partial η^2^ = .01, BF_10_ = 0.21; Repetition Type x Regularity x Attention interaction: *F*(1, 28) = 0.15, *p* = .703, partial η^2^ = .01, BF_10_ = 0.25). Again, amplitudes were overall larger in the attention compared to the no-attention session (main effect of Attention: *F*(1, 28) = 37.86, *p* < .001, partial η^2^ = .57, BF_10_ = 1.57*10^14^), but not significantly influenced by Regularity (main effect of Regularity: *F*(1, 28) = 2.64, *p* = .115, partial η^2^ = .09, BF_10_ = 0.30; Regularity x Attention interaction: *F*(1, 28) = 0.21, *p* = .653, partial η^2^ = .01, BF_10_ = 0.21).

## Discussion

The aim of the present EEG study was to test whether and how temporal regularity of pattern repetitions and listeners’ attention modulate perceptual learning of random acoustic patterns. To this end, we presented participants with correlated noise sequences that contained either temporally regular or jittered repetitions of a certain pattern (manipulation of temporal regularity) while they either performed an auditory repetition detection task or a visual distractor task (manipulation of listeners’ attention towards or away from the pattern repetitions).

We found evidence for the implicit formation of robust memory representations through repeated exposure, reflected in ERP amplitude and behavioural performance differences between reference patterns that recurred across multiple trials and other patterns that were presented in just one trial. Specifically, behavioural sensitivity to pattern repetitions within an auditory sequence increased for the learnt reference patterns compared to other patterns, in line with previous findings with the same or similar types of random acoustic patterns (Agus et al., 2010; Agus & Pressnitzer, 2013, 2021; Bianco et al., 2020; Ringer et al., 2022a; Song & Luo, 2017; Viswanathan et al., 2016). At the electrophysiological level, a negative potential emerged within the first 500 ms relative to the onset of the first pattern presentation and remained sustained throughout the sequence. On top of this sustained potential, pattern-related responses to pattern repetition onsets within a sequence were characterised by a prominent negative deflection between 150 ms and 400 ms (relative to pattern onset), preceded by a weaker positivity. Overall, this combined pattern of sustained potential and pattern-related ERPs, both with a fronto-central distribution, is in line with earlier reports of EEG responses to pattern repetitions embedded in continuous acoustic sequences (Andrillon et al., 2015, 2017; Barascud et al., 2016; Berti et al., 2000; Herrmann et al., 2021; Herrmann & Johnsrude, 2018; Hodapp & Grimm, 2021; Kaernbach et al., 1998; Ringer et al., 2022b). Critically, amplitudes of the pattern-related ERPs and ITPC differed between reference patterns (RefRN) and other patterns (RN): At pattern position two to six within the sequence, ERP amplitudes were less negative between 140 and 220 ms and ITPC was increased within the first 190 ms relative to pattern repetition onset in RefRN compared to RN sequences. These two measures are closely linked and together suggest that memory representations formed through previous exposure to specific reference patterns enhance phase alignment of neural responses to pattern repetitions, which on average across trials leads to respective ERP differences. This pattern of results replicates and brings together previous findings from studies that reported different neural signatures of memory formation for random acoustic patterns: The strengthened phase alignment of pattern-related responses for implicitly memorised patterns is reflected in stronger ITPC (Andrillon et al., 2015; Luo et al., 2013) and a narrower, but larger negativity in the ERP, in turn resulting in decreased amplitudes during the slope (Hodapp & Grimm, 2021) and increased amplitudes at the peak (Andrillon et al., 2015; Andrillon et al., 2017). Strikingly, we also found a significant memory-related difference in the event-related potential already before any within-sequence pattern repetition: The negative amplitude shift was larger for RefRN compared to RN sequences between 260 and 500 ms relative to the first pattern presentation per sequence.

While repeated exposure to the reference pattern across multiple trials increased behavioural repetition detection performance and modified pattern-related ERP amplitudes, memory formation was not reflected in above-chance behavioural recognition of learnt patterns, unlike in a previous study with the same type of stimulus material (Ringer et al., 2022a). Thus, albeit the implicitly acquired memory representations for the reference patterns improved perceptual sensitivity, they could not be actively accessed during the subsequent unexpected memory test (regardless of the manipulations of temporal regularity and listeners’ attention during learning). This discrepancy between more direct and more indirect correlates of memory formation may relate to the fact that active selection of the recognised pattern during the memory test requires more cognitive effort than repetition detection, which rather takes place on a perceptual level. There are several differences in the stimulus design that might explain why above-chance recognition of learnt reference patterns was found in the previous (Ringer et al., 2022a), but not in the current study: In the current study, we presented shorter reference patterns (200 ms in the present vs. 500 ms in the previous study), which decreased the probability that a pattern by chance contained a short characteristic spectral feature that made it easier to memorise. The patterns occurred embedded in a continuous sequence during learning and were presented as short sounds in isolation in the subsequent memory test, whereas there was no such (strong) difference in presentation format between learning and recall in the previous study. Moreover, the number of to-be-learnt reference pattern was larger (10 vs. 3) and the retention interval between implicit learning of a pattern and active recall during the memory test was longer in the current study (up to an hour vs. several minutes). Taken together, all these aspects increased the difficulty of active recognition of previously learnt reference patterns in the unexpected memory test and might have rendered above-chance recognition performance impossible.

### Substantial pattern learning in both regular and jittered sequences

We found that the memory effect was not significantly modulated by the temporal regularity of pattern repetitions. Listeners were able to form memory representations of recurring reference patterns regardless of whether they occurred in regular or jittered auditory sequences. Learning was reflected in a substantial increase of behavioural repetition detection performance and a significant modulation of pattern-related ERPs for both the first and the remaining pattern presentations per sequence (at least in the attention session). No significant difference in magnitude of the memory effect between regularity conditions was found for either of these behavioural and electrophysiological markers. This appears inconsistent with the hypothesis that temporal regularity improves perception (Jones, 1976, 2019; Henry & Herrmann, 2014; Large & Jones, 1999), which in turn might facilitate longer-term memory formation, and with a previous study that reported a reduction of the behavioural memory effect for reference patterns in temporally jittered (compared to isochronous) sequences (Dauer et al., 2022). Conversely, Hodapp & Grimm (2021) found no amplitude modulation of the pattern-related negativity by temporal regularity, just like the current study.

Discrepancies between studies with regard to the effect of temporal regularity on pattern learning might stem from differences in the stimulus material. For instance, due to the correlative structure inherent in correlated compared to white noise, characteristic spectral features (which the randomly generated stimulus contains by chance) are more likely to perceptually “pop out” and facilitate repetition detection. It is plausible to assume that temporal regularity does not enhance perception and memory to the same degree under all circumstances, and that listeners might particularly benefit from rhythmic, predictable stimulus presentation in challenging situations, such as when they are asked to detect repetitions in acoustic material with minimal spectro-temporal structure (e.g., white noise). In contrast, when the task is easier and repetitions are more salient (even in jittered sequences), the facilitation by temporal regularity might become insignificant, such as in the current study.

More generally, there are several possible explanations why temporally regular (compared to jittered) stimulus presentation does not per se improve perceptual pattern learning, which might also account for the absence of a significant modulation of the memory effect by temporal regularity in the current data. One explanation is the extensive experience with temporal jitter in natural sounds during everyday listening situations. Many natural sounds, including environmental sounds, speech, and music, contain temporal regularities at characteristic frequencies, which listeners exploit to sharpen perception (Ding et al., 2017; Doelling & Poeppel, 2015; Giraud & Poeppel, 2012; Poeppel & Assaneo, 2020). However, most of these regularities are not strictly isochronous, but only quasi-periodic (Obleser et al., 2017; Rimmele et al., 2018). Although the brain is naturally tuned to periodic rhythms at different time scales, it exhibits flexibility towards some temporal variability, e.g., cortical tracking was observed also for stimulus rhythms that are not strictly periodic (Breska & Deouell, 2017; Morillon et al., 2016; Obleser et al., 2017; Rimmele et al., 2018). Thus, the human auditory system is highly trained in dealing with temporal jitter in the order of up to a few hundreds of milliseconds, and the jittered condition in our experiment might actually resemble demands of naturalistic listening situations.

Another possibility is that memory formation in jittered blocks primarily happened in trials in which the sequence by chance contained several (adjacent) inter-pattern intervals that were similar in length (or some other rhythm of shorter and longer inter-pattern intervals). We tried to avoid incidentally (almost-) isochronous sequences in the jittered condition by including the restriction that two adjacent inter-pattern intervals must differ in duration by at least 50 ms. However, this cannot fully prevent that the variance in inter-pattern interval duration differs between sequences, resulting in “more jittered” and “less jittered” sequences. Similarly, some sequences contained more shorter inter-pattern intervals (below 300 ms) than others, which, due to less interfering sensory input between consecutive pattern presentations, might also facilitate repetition detection and memory formation. Previous studies reported successful pattern learning from just two pattern presentations per trial (Agus et al., 2010; Agus & Pressnitzer, 2013; Ringer et al., 2022a; Viswanathan et al., 2016) and an amelioration of the disadvantageous effect of temporal jitter when some sequences with isochronous pattern onsets were included in an experimental block (Dauer et al., 2022). Thus, in line with the interpretation by Dauer and colleagues (2022), memories of the reference patterns might have been formed in trials that were incidentally “less jittered” or contained more short inter-pattern intervals and translated to “more jittered” trials. Future studies could clarify whether this hypothesis holds true by systematically manipulating the variation in inter-pattern interval duration.

Taken together, we found evidence for a comparable memory effect across temporally regular and jittered sequences in the current study, suggesting a flexible learning across different degrees of temporal structure. Whether pattern learning relies on the exact same mechanisms in regular and jittered sequences remains to be clarified. We cannot exclude that listeners employed different strategies in regular and jittered blocks, which still resulted in comparable behavioural performance (in the auditory repetition detection task); in fact, the introduction of separate blocks with either regular or irregular repetitions in the instruction might have even encouraged them to tune their perceptual strategy to the demands of the current block. Moreover, it might be plausible that the temporal regularity of pattern repetitions would play a larger role in different (experimental) learning contexts (e.g., with different stimulus material, a different task, or an even larger amount of jitter). That is, the current findings do not rule out the possibility that – at least under certain circumstances – temporal regularity has a significant beneficial impact on memory formation for random acoustic patterns.

### Attention to pattern repetitions boosts learning

Our data provided evidence that attention to the pattern repetitions improved perceptual learning of random acoustic patterns. We found that the memory effect was larger at the first pattern position per sequence when participants directed their attention towards the auditory stimulation compared to when they performed a visual distractor task. Thus, enhanced perceptual representations through an attentional focus on the auditory stimulation supported the formation of longer-term memories for initially unfamiliar acoustic patterns. In particular, attention might have facilitated access to established memory representations for previously presented patterns upon first occurrence with a sequence.

It is important to note that, despite the modulation of the memory effect by attention, perceptual learning was still possible during inattention. This is in line with previous studies that reported a significant memory effect during an auditory distractor task (Andrillon et al., 2015) or REM sleep (Andrillon et al., 2017). Clear evidence for a modulation of the memory effect by attention was observed for the first pattern presentation within a sequence in the current data, which suggests that listeners’ attentional state specifically affects certain aspects of learning and recognition of random acoustic patterns. We suspect that the reduction of the memory effect might have resulted from a delayed onset of the effect within the block in the absence of attention, which consequently led to a diminished memory effect when averaging over the whole block. To test this hypothesis, we conducted a supplementary analysis and compared pattern-related ERP amplitudes at the first pattern position per sequence between reference and other patterns separately for an early trial group (i.e., when not much learning should have happened yet) and a late trial group (i.e., after learning could have happened) within the block (see Supplemental Material). This analysis showed a similar increase of the memory effect from the beginning to the end of the block across all regularity and attention conditions. These findings support our idea that the difference between attention conditions does not arise from a difference in magnitude of the memory effect after learning has taken place, but likely from a speed-up of learning through attention to the pattern repetitions.

The fixed session order (i.e., no-attention first, attention second) can be considered a shortcoming of our experimental design. We chose this procedure to avoid that participants were aware of the pattern repetitions in the auditory stimulation during the no-attention session after performing the repetition detection in the session before. However, one might argue that the enlarged memory effect in the attention session cannot only be attributed to the listeners’ attentional focus on the auditory pattern repetitions (instead of away from them), but could also stem from increased (passive) familiarisation with the stimulus material during the preceding no-attention session. Nevertheless, we argue that this methodological limitation does not question the interpretability of the attention effect: While passive exposure to the stimulus material might have improved overall repetition detection, it is unlikely that it explains the item-specific memory effect for particular reference patterns, since we used different reference patterns in both sessions and previous studies showed striking differences in learning success between blocks (Agus et al., 2010), suggesting no straightforward relationship between experience with the stimulus material and memory formation for single specific patterns.

Taken together, although inattention does not preclude perceptual learning of random acoustic patterns, memory formation appears to benefit from attention to pattern repetitions in the auditory stimulation. This finding extends previous research by directly contrasting memory formation with and without attention, whereas earlier studies only focussed on either condition in isolation.

### Memory effect is most pronounced at the beginning of the sequence

We found that the memory effect was most pronounced at the first pattern position early in the auditory sequence. This time window is particularly critical for assessing longer-term learning because before the first within-sequence pattern repetition any effect should be related to memory formation across trials. Only few studies investigated neural correlates of perceptual learning at the first pattern presentation per sequence, and only few studies statistically analysed the first and later pattern positions separately. Hodapp & Grimm (2021) found no evidence for a memory effect at the first position, whereas there was a significant effect at later positions (Hodapp & Grimm, 2021).

Conversely, Andrillon and colleagues (2015) did report a significant cluster of ERP difference between the learnt reference and other patterns already after the first pattern onset (Andrillon et al., 2015). Similarly to our current data, visual inspection of EEG responses to the whole sequence suggests that the memory effect in pattern-related ERP amplitude tended to decrease at later pattern positions throughout the sequence (Andrillon et al., 2015). A plausible explanation for this observation is that sequences that contain the learnt reference pattern differ most from sequences that contain repetitions of a novel pattern right at the beginning: Only the occurrence of the reference pattern allows to anticipate upcoming pattern repetitions (since the reference pattern always occurs in sequences that contain repetitions) based on memory representations formed in previous trials, while repetitions of a novel pattern can be detected at the earliest at the second pattern position. Throughout the remaining sequence, repetitions become salient and distinguishable from sequences without repetitions irrespective of whether the reference or another pattern is repeated. Previous findings (e.g., Barascud et al., 2016) suggest that repetitions are detected as early as during the second pattern occurrence, which was likely also the case in the present study (just the response was delayed via the task instruction for the repetition detection task to avoid contamination of the auditory ERPs with motor activity). Furthermore, this early phase of the sequence appears to be most susceptible to a processing benefit through attention, which might facilitate (implicit) recognition of the reference pattern upon its first occurrence.

## Conclusions

The current study sought to scrutinise a potential modulation of perceptual learning of random acoustic patterns by temporal regularity of pattern repetitions and listeners’ attention. Our results support previous findings of implicit memory formation for specific, initially unfamiliar acoustic patterns through repeated exposure. Furthermore, we showed that the learning mechanism is flexible and allows to successfully build up memory representations across both temporally regular and jittered sound sequences. While learning does not necessarily require attention to the repetitions in the stimulation, attentional focus benefits, i.e., speeds up, memory formation and, in particular, facilitates the access to established memory representations upon first occurrence within a sequence. These findings both highlight the flexibility of perceptual learning mechanisms across naturally varying auditory environments and provide insights about how features of the learning context can boost memory formation.

## Supporting information

supplementary analysis

## Acknowledgements

The authors thank Benjamin Eichenberger for help with EEG data collection.

## Data Availability

Data of individual participants and scripts to reproduce the statistical analysis reported in the manuscript can be found here: https://osf.io/r8gea/?view_only=4580e185befa474eac7a2d0efb3c615e

Further data and materials are available from the corresponding author upon reasonable request.

## References

Agus, T. R., & Pressnitzer, D. (2013). The detection of repetitions in noise before and after perceptual learning. The Journal of the Acoustical Society of America, 134(1), 464–473. https://doi.org/10.1121/1.4807641

Agus, T. R., & Pressnitzer, D. (2021). Repetition detection and rapid auditory learning for stochastic tone clouds. The Journal of the Acoustical Society of America, 150(3), 1735–1749. https://doi.org/10.1121/10.0005935

Agus, T. R., Thorpe, S. J., & Pressnitzer, D. (2010). Rapid Formation of Robust Auditory Memories: Insights from Noise. Neuron, 66(4), 610–618. https://doi.org/10.1016/j.neuron.2010.04.014

Andreou, L.-V., Kashino, M., & Chait, M. (2011). The role of temporal regularity in auditory segregation. Hearing Research, 280(1–2), 228–235. https://doi.org/10.1016/j.heares.2011.06.001

Andrillon, T., Kouider, S., Agus, T., & Pressnitzer, D. (2015). Perceptual Learning of Acoustic Noise Generates Memory-Evoked Potentials. Current Biology, 25(21), 2823–2829. https://doi.org/10.1016/j.cub.2015.09.027

Andrillon, T., Pressnitzer, D., Léger, D., & Kouider, S. (2017). Formation and suppression of acoustic memories during human sleep. Nature Communications, 8(1), 179. https://doi.org/10.1038/s41467-017-00071-z

Banai, K., & Lavie, L. (2020). Rapid Perceptual Learning and Individual Differences in Speech Perception: The Good, the Bad, and the Sad. Auditory Perception & Cognition, 3(4), 201–211. https://doi.org/10.1080/25742442.2021.1909400

Barascud, N., Pearce, M. T., Griffiths, T. D., Friston, K. J., & Chait, M. (2016). Brain responses in humans reveal ideal observer-like sensitivity to complex acoustic patterns. Proceedings of the National Academy of Sciences, 113(5), E616–E625. https://doi.org/10.1073/pnas.1508523113

Barnes, R., & Jones, M. R. (2000). Expectancy, Attention, and Time. Cognitive Psychology, 41(3), 254–311. https://doi.org/10.1006/cogp.2000.0738

Bergh, D. van den, Doorn, J. van, Marsman, M., Draws, T., Kesteren, E.-J. van, Derks, K., Dablander, F., Gronau, Q. F., Kucharský, Š., Gupta, A. R. K. N., Sarafoglou, A., Voelkel, J. G., Stefan, A., Ly, A., Hinne, M., Matzke, D., & Wagenmakers, E.-J. (2020). A tutorial on conducting and interpreting a Bayesian ANOVA in JASP. LAnnee Psychologique, 120(1), 73–96. https://www.cairn-int.info/journal--2020-1-page-73.htm

Berti, S., Schröger, E., & Mecklinger, A. (2000). Attentive and pre-attentive periodicity analysis in auditory memory: An event-related brain potential study. Neuroreport, 11(9), 1883–1887. https://doi.org/10.1097/00001756-200006260-00016

Bianco, R., Harrison, P. M., Hu, M., Bolger, C., Picken, S., Pearce, M. T., & Chait, M. (2020). Long-term implicit memory for sequential auditory patterns in humans. ELife, 9, e56073. https://doi.org/10.7554/eLife.56073

Brainard, D. H. (1997). The psychophysics toolbox. Spatial Vision, 10(4), 433–436. https://doi.org/10.1163/156856897X00357

Bregman, A. S. (1990). Auditory scene analysis: The perceptual organization of sound. MIT Press.

Breska, A., & Deouell, L. Y. (2017). Neural mechanisms of rhythm-based temporal prediction: Delta phase-locking reflects temporal predictability but not rhythmic entrainment. PLOS Biology, 15(2), e2001665. https://doi.org/10.1371/journal.pbio.2001665

Calderone, D. J., Lakatos, P., Butler, P. D., & Castellanos, F. X. (2014). Entrainment of neural oscillations as a modifiable substrate of attention. Trends in Cognitive Sciences, 18(6), 300–309. https://doi.org/10.1016/j.tics.2014.02.005

Chang, A., Bosnyak, D. J., & Trainor, L. J. (2019). Rhythmicity facilitates pitch discrimination: Differential roles of low and high frequency neural oscillations. NeuroImage, 198, 31–43. https://doi.org/10.1016/j.neuroimage.2019.05.007

Chun, M. M., & Turk-Browne, N. B. (2007). Interactions between attention and memory. Current Opinion in Neurobiology, 17(2), 177–184. https://doi.org/10.1016/j.conb.2007.03.005

Dauer, T., Henry, M. J., & Herrmann, B. (2022). Auditory perceptual learning depends on temporal regularity and certainty. Journal of Experimental Psychology: Human Perception and Performance, 48(7), 755–770. https://doi.org/10.1037/xhp0001016

Delorme, A., & Makeig, S. (2004). EEGLAB: An open source toolbox for analysis of single-trial EEG dynamics including independent component analysis. Journal of Neuroscience Methods, 134(1), 9–21. https://doi.org/10.1016/j.jneumeth.2003.10.009

Ding, N., Patel, A. D., Chen, L., Butler, H., Luo, C., & Poeppel, D. (2017). Temporal modulations in speech and music. Neuroscience & Biobehavioral Reviews, 81, 181–187. https://doi.org/10.1016/j.neubiorev.2017.02.011

Doelling, K. B., & Poeppel, D. (2015). Cortical entrainment to music and its modulation by expertise. Proceedings of the National Academy of Sciences, 112(45), E6233–E6242. https://doi.org/10.1073/pnas.1508431112

Eaton, J. W., Bateman, D., Hauberg, S., & Wehbring, R. (2019). GNU Octave version 5.2.0 manual: a high-level interactive language for numerical comparisons. https://www.gnu.org/software/octave/doc/v5.2.0/

Geiser, E., Notter, M., & Gabrieli, J. D. E. (2012). A Corticostriatal Neural System Enhances Auditory Perception through Temporal Context Processing. Journal of Neuroscience, 32(18), 6177– 6182. https://doi.org/10.1523/JNEUROSCI.5153-11.2012

Gibson, E. J. (1969). Principles of perceptual learning and development. Appleton-Century-Crofts.

Gilbert, C. D., Sigman, M., & Crist, R. E. (2001). The Neural Basis of Perceptual Learning. Neuron, 31(5), 681–697. https://doi.org/10.1016/S0896-6273(01)00424-X

Giraud, A.-L., & Poeppel, D. (2012). Cortical oscillations and speech processing: Emerging computational principles and operations. Nature Neuroscience, 15(4), 511–517. https://doi.org/10.1038/nn.3063

Hanslmayr, S., Axmacher, N., & Inman, C. S. (2019). Modulating Human Memory via Entrainment of Brain Oscillations. Trends in Neurosciences, 42(7), 485–499. https://doi.org/10.1016/j.tins.2019.04.004

Hautus, M. J., & Lee, A. (2006). Estimating sensitivity and bias in a yes/no task. British Journal of Mathematical and Statistical Psychology, 59(2), 257–273. https://doi.org/10.1348/000711005X65753

Henry, M. J., & Herrmann, B. (2014). Low-Frequency Neural Oscillations Support Dynamic Attending in Temporal Context. Timing & Time Perception, 2(1), 62–86. https://doi.org/10.1163/22134468-00002011

Henry, M. J., & Obleser, J. (2012). Frequency modulation entrains slow neural oscillations and optimizes human listening behavior. Proceedings of the National Academy of Sciences, 109(49), 20095–20100. https://doi.org/10.1073/pnas.1213390109

Herrmann, B., Araz, K., & Johnsrude, I. S. (2021). Sustained neural activity correlates with rapid perceptual learning of auditory patterns. NeuroImage, 238, 118238. https://doi.org/10.1016/j.neuroimage.2021.118238

Herrmann, B., & Johnsrude, I. S. (2018). Neural Signatures of the Processing of Temporal Patterns in Sound. The Journal of Neuroscience, 38(24), 5466–5477. https://doi.org/10.1523/JNEUROSCI.0346-18.2018

Hickey, P., & Race, E. (2021). Riding the slow wave: Exploring the role of entrained low-frequency oscillations in memory formation. Neuropsychologia, 160, 107962. https://doi.org/10.1016/j.neuropsychologia.2021.107962

Hodapp, A., & Grimm, S. (2021). Neural signatures of temporal regularity and recurring patterns in random tonal sound sequences. European Journal of Neuroscience, 53(8), 2740–2754. https://doi.org/10.1111/ejn.15123

Huyck, J. J., & Johnsrude, I. S. (2012). Rapid perceptual learning of noise-vocoded speech requires attention. The Journal of the Acoustical Society of America, 131(3), EL236–EL242. https://doi.org/10.1121/1.3685511

Irvine, D. R. F. (2018). Auditory perceptual learning and changes in the conceptualization of auditory cortex. Hearing Research, 366, 3–16. https://doi.org/10.1016/j.heares.2018.03.011

Irvine, D. R. F., Martin, R. L., Klimkeit, E., & Smith, R. (2000). Specificity of perceptual learning in a frequency discrimination task. The Journal of the Acoustical Society of America, 108(6), 2964–2968. https://doi.org/10.1121/1.1323465

Jones, M. R. (1976). Time, our lost dimension: Toward a new theory of perception, attention, and memory. Psychological Review, 83, 323–355. https://doi.org/10.1037/0033-295X.83.5.323

Jones, M. R., Johnston, H. M., & Puente, J. (2006). Effects of auditory pattern structure on anticipatory and reactive attending. Cognitive Psychology, 53(1), 59–96. https://doi.org/10.1016/j.cogpsych.2006.01.003

Jones, M. R., Moynihan, H., MacKenzie, N., & Puente, J. (2002). Temporal Aspects of Stimulus-Driven Attending in Dynamic Arrays. Psychological Science, 14(4), 313–319. https://doi.org/10.1111/1467-9280.00458

Kaernbach, C., Schröger, E., & Gunter, T. C. (1998). Human event-related brain potentials to auditory periodic noise stimuli. Neuroscience Letters, 242(1), 17–20. https://doi.org/10.1016/S0304-3940(98)00034-2

Kang, H., Agus, T. R., & Pressnitzer, D. (2017). Auditory memory for random time patterns. The Journal of the Acoustical Society of America, 142(4), 2219–2232. https://doi.org/10.1121/1.5007730

Kang, H., Lancelin, D., & Pressnitzer, D. (2018). Memory for Random Time Patterns in Audition, Touch, and Vision. Neuroscience, 389, 118–132. https://doi.org/10.1016/j.neuroscience.2018.03.017

Kang, H., Macherey, O., Roman, S., & Pressnitzer, D. (2021). Auditory memory for random time patterns in cochlear implant listeners. The Journal of the Acoustical Society of America, 150(3), 1934–1944. https://doi.org/10.1121/10.0005728

Kleiner, M., Brainard, D., Pelli, D. G., Ingling, A., Murray, R., & Broussard, C. (2007). “What’s new in Psychtoolbox-3?”. Perception 36 ECVP Abstract Supplement.

Kraus, N., & Banai, K. (2007). Auditory-Processing Malleability: Focus on Language and Music. Current Directions in Psychological Science, 16(2), 105–110. https://doi.org/10.1111/j.1467-8721.2007.00485.x

Kumar, S., Bonnici, H. M., Teki, S., Agus, T. R., Pressnitzer, D., Maguire, E. A., & Griffiths, T. D. (2014). Representations of specific acoustic patterns in the auditory cortex and hippocampus. Proceedings of the Royal Society B: Biological Sciences, 281(1791), 20141000. https://doi.org/10.1098/rspb.2014.1000

Lakatos, P., Gross, J., & Thut, G. (2019). A New Unifying Account of the Roles of Neuronal Entrainment. Current Biology, 29(18), R890–R905. https://doi.org/10.1016/j.cub.2019.07.075

Lakatos, P., Karmos, G., Mehta, A. D., Ulbert, I., & Schroeder, C. E. (2008). Entrainment of Neuronal Oscillations as a Mechanism of Attentional Selection. Science, 320(5872), 110–113. https://doi.org/10.1126/science.1154735

Lakatos, P., O’Connell, M. N., Barczak, A., Mills, A., Javitt, D. C., & Schroeder, C. E. (2009). The Leading Sense: Supramodal Control of Neurophysiological Context by Attention. Neuron, 64(3), 419–430. https://doi.org/10.1016/j.neuron.2009.10.014

Lakatos, P., Shah, A. S., Knuth, K. H., Ulbert, I., Karmos, G., & Schroeder, C. E. (2005). An Oscillatory Hierarchy Controlling Neuronal Excitability and Stimulus Processing in the Auditory Cortex. Journal of Neurophysiology, 94(3), 1904–1911. https://doi.org/10.1152/jn.00263.2005

Large, E. W., & Jones, M. R. (1999). The dynamics of attending: How people track time-varying events. Psychological Review, 106(1), 119–159. https://doi.org/10.1037/0033-295X.106.1.119

Lawrance, E. L. A., Harper, N. S., Cooke, J. E., & Schnupp, J. W. H. (2014). Temporal predictability enhances auditory detection. The Journal of the Acoustical Society of America, 135(6), EL357–EL363. https://doi.org/10.1121/1.4879667

Lawrence, M. A. (2016). ez: Easy analysis and visualization of factorial experiments. R package version 4.4-0. https://CRAN.Rproject.org/packa ge=ez

Lee, M. D., & Wagenmakers, E.-J. (2014). Bayesian Cognitive Modeling: A Practical Course. Cambridge University Press.

Luck, S. J., & Gaspelin, N. (2017). How to get statistically significant effects in any ERP experiment (and why you shouldn’t). Psychophysiology, 54(1), 146–157. https://doi.org/10.1111/psyp.12639

Luo, H., Tian, X., Song, K., Zhou, K., & Poeppel, D. (2013). Neural Response Phase Tracks How Listeners Learn New Acoustic Representations. Current Biology, 23(11), 968–974. https://doi.org/10.1016/j.cub.2013.04.031

Macmillan, N. A. (2001). Signal Detection Theory. In N. J. Smelser & P. B. Baltes (Eds.), International Encyclopedia of the Social & Behavioral Sciences (pp. 14075–14078). Pergamon. https://doi.org/10.1016/B0-08-043076-7/00677-X

Maris, E. (2012). Statistical testing in electrophysiological studies. Psychophysiology, 49(4), 549– 565.https://doi.org/10.1111/j.1469-8986.2011.01320.x

Maris, E., & Oostenveld, R. (2007). Nonparametric statistical testing of EEG-and MEG-data. Journal of Neuroscience Methods, 164(1), 177–190. https://doi.org/10.1016/j.jneumeth.2007.03.024

Masutomi, K., Barascud, N., Kashino, M., McDermott, J. H., & Chait, M. (2016). Sound segregation via embedded repetition is robust to inattention. Journal of Experimental Psychology: Human Perception and Performance, 42(3), 386–400. https://doi.org/10.1037/xhp0000147

McDermott, J. H., Schemitsch, M., & Simoncelli, E. P. (2013). Summary statistics in auditory perception. Nature Neuroscience, 16(4), 493–498. https://doi.org/10.1038/nn.3347

McDermott, J. H., Wrobleski, D., & Oxenham, A. J. (2011). Recovering sound sources from embedded repetition. Proceedings of the National Academy of Sciences, 108(3), 1188–1193. https://doi.org/10.1073/pnas.1004765108

Morey, R. D., & Rouder, J. N. (2011). Bayes factor approaches for testing interval null hypotheses. Psychological Methods, 16(4), 406–419. https://doi.org/10.1037/a0024377

Morey, R. D., Rouder, J. N., Jamil, T., Urbanek, S., Forner, K., & Ly, A. (2018). BayesFactor: Computation of Bayes Factors for common designs. R package version 0.9.12-4.2. https://cran.r-project.org/package=BayesFactor

Morillon, B., Schroeder, C. E., Wyart, V., & Arnal, L. H. (2016). Temporal Prediction in lieu of Periodic Stimulation. The Journal of Neuroscience, 36(8), 2342–2347. https://doi.org/10.1523/JNEUROSCI.0836-15.2016

Norris, D., McQueen, J. M., & Cutler, A. (2003). Perceptual learning in speech. Cognitive Psychology, 47(2), 204–238. https://doi.org/10.1016/S0010-0285(03)00006-9

Obleser, J., Henry, M. J., & Lakatos, P. (2017). What do we talk about when we talk about rhythm? PLOS Biology, 15(9), e2002794. https://doi.org/10.1371/journal.pbio.2002794

Oldfield, R. C. (1971). The Edinburgh handedness inventory. Neuropsychologia, 9, 97–111. https://doi.org/10.1002/mus.10529

Oostenveld, R., Fries, P., Maris, E., & Schoffelen, J. M. (2011). FieldTrip: Open source software for advanced analysis of MEG, EEG, and invasive electrophysiological data. Computational Intelligence and Neuroscience, 2011, 1–9. https://doi.org/10.1155/2011/156869

Picton, T. W., Woods, D. L., & Proulx, G. B. (1978). Human auditory sustained potentials. I. The nature of the response. Electroencephalography and Clinical Neurophysiology, 45(2), 186–197. https://doi.org/10.1016/0013-4694(78)90003-2

Pion-Tonachini, L., Kreutz-Delgado, K., & Makeig, S. (2019). ICLabel: An automated electroencephalographic independent component classifier, dataset, and website. NeuroImage, 198, 181–197. https://doi.org/10.1016/j.neuroimage.2019.05.026

Poeppel, D., & Assaneo, M. F. (2020). Speech rhythms and their neural foundations. Nature Reviews Neuroscience, 21(6), 322–334. https://doi.org/10.1038/s41583-020-0304-4

Rajendran, V. G., Harper, N. S., Abdel-Latif, K. H. A., & Schnupp, J. W. H. (2016). Rhythm Facilitates the Detection of Repeating Sound Patterns. Frontiers in Neuroscience, 10. https://doi.org/10.3389/fnins.2016.00009

Jones, M. R. (2019). Time Will Tell: A Theory of Dynamic Attending. Oxford University Press. https://doi.org/10.1093/oso/9780190618216.001.0001

Rimmele, J. M., Morillon, B., Poeppel, D., & Arnal, L. H. (2018). Proactive Sensing of Periodic and Aperiodic Auditory Patterns. Trends in Cognitive Sciences, 22(10), 870–882. https://doi.org/10.1016/j.tics.2018.08.003

Ringer, H., Schröger, E., & Grimm, S. (2022a). Perceptual Learning and Recognition of Random Acoustic Patterns. Auditory Perception & Cognition, 0(0), 1–23. https://doi.org/10.1080/25742442.2022.2082827

Ringer, H., Schröger, E., & Grimm, S. (2022b). Within- and between-subject consistency of perceptual segmentation in periodic noise: A combined behavioral tapping and EEG study. *Psychophysiology*, e14174. https://doi.org/10.1111/psyp.14174

Rouder, J. N., Morey, R. D., Speckman, P. L., & Province, J. M. (2012). Default Bayes factors for ANOVA designs. Journal of Mathematical Psychology, 56(5), 356–374. https://doi.org/10.1016/j.jmp.2012.08.001

Rouder, J. N., Speckman, P. L., Sun, D., Morey, R. D., & Iverson, G. (2009). Bayesian t tests for accepting and rejecting the null hypothesis. Psychonomic Bulletin & Review, 16(2), 225–237. https://doi.org/10.3758/PBR.16.2.225

Samuel, A. G., & Kraljic, T. (2009). Perceptual learning for speech. Attention, Perception, & Psychophysics, 71(6), 1207–1218. https://doi.org/10.3758/APP.71.6.1207

Schroeder, C. E., & Lakatos, P. (2009). Low-frequency neuronal oscillations as instruments of sensory selection. Trends in Neurosciences, 32(1), 9–18. https://doi.org/10.1016/j.tins.2008.09.012

Sohoglu, E., & Chait, M. (2016). Detecting and representing predictable structure during auditory scene analysis. ELife, 5, e19113. https://doi.org/10.7554/eLife.19113

Song, K., & Luo, H. (2017). Temporal Organization of Sound Information in Auditory Memory. Frontiers in Psychology, 8, 999. https://doi.org/10.3389/fpsyg.2017.00999

Southwell, R., Baumann, A., Gal, C., Barascud, N., Friston, K., & Chait, M. (2017). Is predictability salient? A study of attentional capture by auditory patterns. Philosophical Transactions of the Royal Society B: Biological Sciences, 372(1714), 20160105. https://doi.org/10.1098/rstb.2016.0105

Southwell, R., & Chait, M. (2018). Enhanced deviant responses in patterned relative to random sound sequences. Cortex, 109, 92–103. https://doi.org/10.1016/j.cortex.2018.08.032

Viswanathan, J., Rémy, F., Bacon-Macé, N., & Thorpe, S. J. (2016). Long Term Memory for Noise: Evidence of Robust Encoding of Very Short Temporal Acoustic Patterns. Frontiers in Neuroscience, 10. https://doi.org/10.3389/fnins.2016.00490

Wild, C. J., Yusuf, A., Wilson, D. E., Peelle, J. E., Davis, M. H., & Johnsrude, I. S. (2012). Effortful Listening: The Processing of Degraded Speech Depends Critically on Attention. Journal of Neuroscience, 32(40), 14010–14021. https://doi.org/10.1523/JNEUROSCI.1528-12.2012

Winkler, I., Denham, S. L., & Nelken, I. (2009). Modeling the auditory scene: Predictive regularity representations and perceptual objects. Trends in Cognitive Sciences, 13(12), 532–540. https://doi.org/10.1016/j.tics.2009.09.003

Woods, K. J. P., & McDermott, J. H. (2018). Schema learning for the cocktail party problem. Proceedings of the National Academy of Sciences, 115(14), E3313–E3322. https://doi.org/10.1073/pnas.1801614115

Wright, B. A., & Zhang, Y. (2009). Insights into human auditory processing gained from perceptual learning. In The cognitive neurosciences, 4th ed (pp. 353–365). Massachusetts Institute of Technology.

