## supplementary analysis for "Perceptual learning of random acoustic patterns: Impact of temporal regularity and attention"

### Supplemental Material

When averaging mean RefRN and RN amplitudes over the whole experimental block, this includes trials early in the block in which, after just a few occurrences, no learning of the reference pattern has taken place yet. As we do not expect a difference between the two conditions before learning, this inevitably results in a reduced memory effect overall, defined as the average difference between RefRN and RN. While this does not have an impact as long as learning happens after a similar number of RefRN trials across Regularity and Attention conditions, it might bias the comparison between conditions when the memory benefit for RefRN over RN emerges at rather different time points within the block. Specifically, a later (vs. earlier) onset of learning or slower (vs. steeper) learning trajectory would result in a diminished memory effect over the whole block, although the difference between RefRN and RN might reach a comparable magnitude towards the end of the block. It may be plausible to assume that a concurrent demanding visual distractor task and/or temporal irregularity make the formation of robust memories for the reference pattern more challenging, thereby delaying the learning process. For instance, a similar effect was recently demonstrated for patterns that were separated by a temporal delay or a masker sound between presentations compared to patterns that were presented back-to-back (Ringer et al., 2022).

In a supplemental analysis, we took a closer look at separate early and late portions of the blocks, thus reducing the potential influence of diverging learning trajectories between conditions. That way, we could test whether the modulation of the memory effect by attention, which we found at the first pattern position, was driven by an actual difference in magnitude specifically at the end of the block. Moreover, this analysis allowed to demonstrate that the difference between RefRN and

RN emerged from the beginning to the end of the block, indicating that it actually arises as a result of repeated exposure to the reference pattern in RefRN trials. We chose to contrast the first five RefRN and RN trials per block with the last five RefRN and RN trials, excluding the middle five trials during which most of the behavioural performance change took place in earlier studies (Agus et al., 2010; Ringer et al., 2022). As the memory effect was only modulated by attention at the first pattern presentation within the sequence, we focused on this position to compare mean amplitudes between early and late portions of the blocks.

**Supplemental Methods**

The analysis followed the same procedure as the main analysis of pattern-related responses to the first pattern presentation, with the only exception that RefRN and RN epochs were averaged separately for the early (trial 1-5) and late (trial 11-15) group of trials per block in each condition. Mean amplitudes were extracted from the same time window (260 to 500 ms relative to pattern onset) at electrode Fz. We computed a four-way repeated-measures ANOVA to compare mean amplitudes with the additional factor Trial Group (early, late) beyond Repetition Type, Regularity and Attention. Where applicable, a correction for non-sphericity was used as described for the main analysis, and we again computed both frequentist and Bayesian tests. We expected a significant interaction of Repetition Type and Trial Group, reflecting an increase in magnitude of the memory effect throughout the blocks. Specifically, we hypothesised that RefRN and RN amplitudes differ clearly in the late trial group after learning of the reference pattern has taken place, while there is no (or only a small) difference in the early trial group (in which no or only very little learning has occurred yet). Any three-way or four-way interaction of Repetition Type x Trial Group with Regularity and/or attention would indicate that the increase in magnitude of the memory effect throughout the block is modulated by the respective factor.

**Supplemental Results**

As expected, we found significantly larger negative amplitudes for RefRN than for RN sequences (main effect of Repetition Type: $F(1, 28) = 7.74$, $p = .010$, partial $\eta^2 = .22$, $BF_{10} = 6.51$) and a significant interaction between Repetition Type and Trial Group ($F(1, 28) = 4.95$, $p = .034$, partial $\eta^2 = .15$, $BF_{10} = 2.98$), which indicated that the memory effect increased from the beginning to the end of the block. Importantly, this increase of the memory effect throughout the block was not further modulated by Regularity and Attention (three-way and four-way interactions involving Repetition Type and Trial Group: all $p$'s > .249, all $BF_{10}$'s < 0.21). This pattern of results suggests that the enlarged memory effect in the attention compared to the no-attention session that we observed over the whole block in the main analysis does likely not reflect a larger magnitude of the memory effect after learning has taken place (i.e., at the end of the block). Instead, it is plausible that attention to the acoustic pattern repetitions speeds up learning, which results in a larger memory effect when averaging across the whole block.

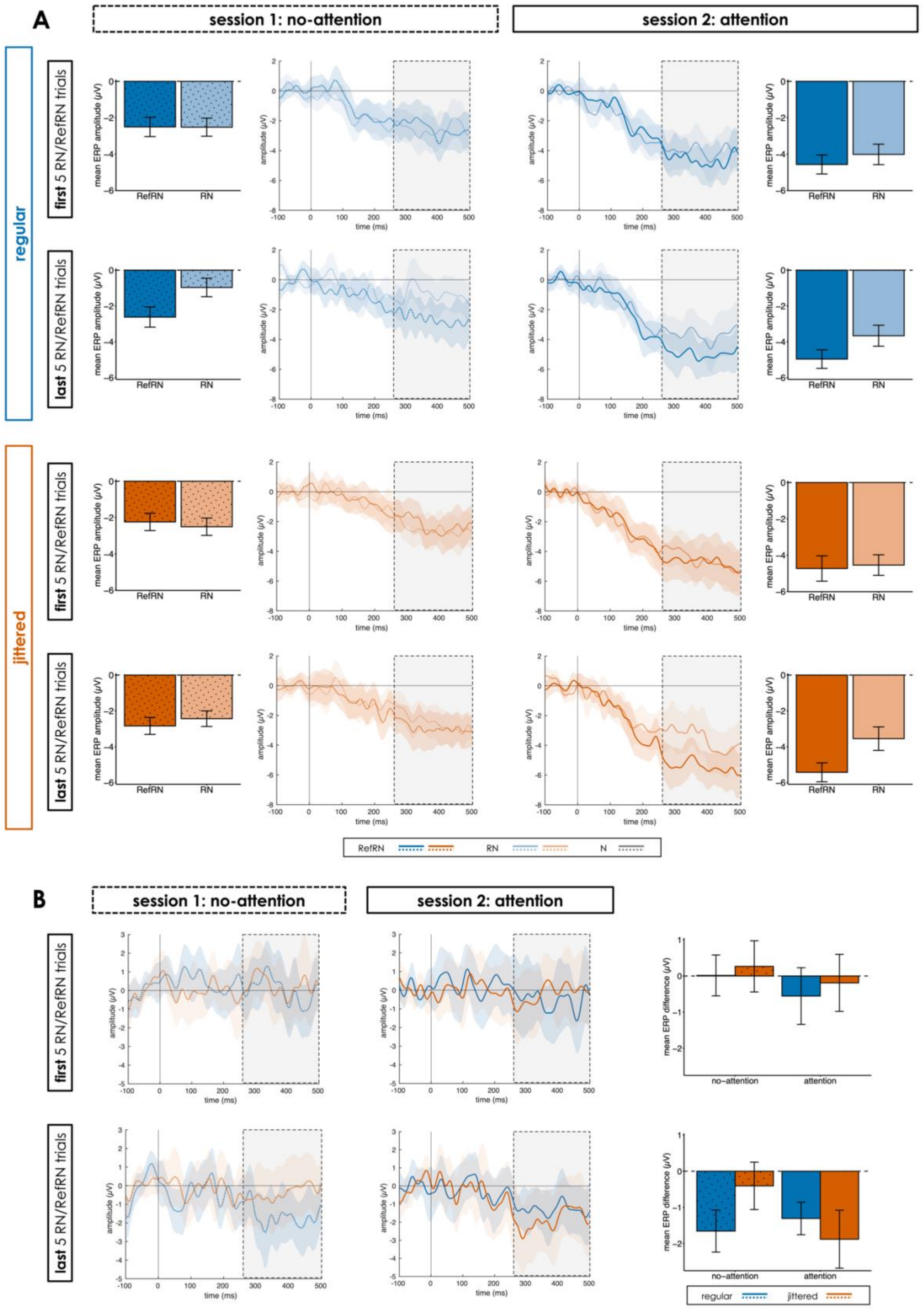
A
session 1: no-attention
session 2: attention
regular
jittered
first 5 RN/RefRN trials
last 5 RN/RefRN trials
mean ERP amplitude (μV)
RefRN
RN
amplitude (μV)
time (ms)
RefRN
RN
N
B
session 1: no-attention
session 2: attention
first 5 RN/RefRN trials
last 5 RN/RefRN trials
amplitude (μV)
time (ms)
mean ERP difference (μV)
no-attention
attention
regular
jittered

*Figure S1.* A: Middle panels: Pattern-related response relative to pattern onset at the first pattern position within the sequence (0 ms) at electrode Fz for N, RN and RefRN sequences in the four Regularity (regular, jittered) and Attention (attention, no-attention) conditions, separately for an early (trial 1-5) and a late (trial 11-15) trial group. Outer panels: Mean amplitudes in the time window of interest (260 to 500 ms relative to the first pattern onset) for RefRN and RN sequences. B: Left panel: Difference waveforms (RefRN-minus-RN) for the four Regularity and Attention conditions in the early and late trial group. Right panel: Mean amplitudes of the difference waveforms. Shaded areas in the ERP plots and error bars in the bar plots indicate ± 1 SEM.

**References**

Agus, T. R., Thorpe, S. J., & Pressnitzer, D. (2010). Rapid Formation of Robust Auditory Memories: Insights from Noise. *Neuron*, *66*(4), 610–618. https://doi.org/10.1016/j.neuron.2010.04.014

Ringer, H., Schröger, E., & Grimm, S. (2022a). Perceptual Learning and Recognition of Random Acoustic Patterns. *Auditory Perception & Cognition*, *0*(0), 1–23. https://doi.org/10.1080/25742442.2022.2082827
